# Antibodies blocking PlGF or VEGF interactions with the NRP1 receptor mediate anti-proliferative effects

**DOI:** 10.1101/2025.10.25.684565

**Authors:** Samuel A. Blackman, Ahlam N. Qerqez, Alison G. Lee, Nicole V. Johnson, Justin M. Owens, Grace S. Lai, Emma C. Aldrich, Kayla G. Sprenger, Jangsoon Lee, Sophie Verrier, Martin J. Stoddart, Annalee W. Nguyen, Jennifer A. Maynard

## Abstract

Antibodies blocking the function of vascular endothelial growth factor A (VEGFA) remain a promising therapeutic strategy, especially when combined with check-point inhibitors, but their efficacy is limited by tumor resistance. This can occur via multiple mechanisms, including upregulation of placental growth factor 2 (PlGF-2), an alternative ligand for VEGF receptor 1 (VEGFR1) and neuropilin receptor 1 (NRP1). Activity of both growth factors is mediated by interactions with multiple receptors and extra-cellular matrix components, which complicates efforts to understand their contributions to cancer progression. To complement existing antibodies, we discovered those blocking interactions between PlGF-2 or VEGFA and their shared NRP1 receptor in the presence of heparin. Limiting angiogenesis to promote vascular normalization is one mechanism of anti-VEGF protection; here, anti-VEGFA antibodies blocking interactions with VEGFR1 and NRP1 reduced HUVEC tube formation in a physiological angiogenesis model. By contrast, antibodies binding PlGF-2 or VEGFA to block NRP1 significantly reduced proliferation of Caki-I kidney carcinoma cells *in vitro*, indicating this receptor mediates additional effects. Interestingly, one antibody exhibited dual-reactive binding to VEGFA and PlGF-2, suggesting a novel therapeutic strategy to prevent PlGF-driven VEGF-resistance. Overall, these antibodies define new mechanisms to disrupt PlGF activity and support a role for NRP1 in cell proliferation.

**Key Results:**

- New antibodies binding VEGFA or PlGF selectively block NRP1-receptor interactions.
- Antibody blockade of VEGFA binding to NRP1 reduced HUVEC angiogenesis.
- Blocking growth factor interactions with NRP1 reduced Caki-I renal carcinoma proliferation.
- Identified an antibody with dual-reactive binding to VEGFA and PlGF.

## Introduction

Two decades after their debut as cancer therapeutics, antibodies that inhibit angiogenesis are generating renewed interest. Recent clinical trials combining the FDA-approved bevacizumab with immune checkpoint inhibitors have demonstrated improved patient survival, acceptable safety profiles, and enhanced immune cell infiltration in melanoma and pancreatic cancer [1–3]. While most anti-angiogenesis candidates have targeted the prototypical vascular endothelial growth factor variant A (VEGFA) to normalize tumor vasculature and thereby enhance tumor exposure to cytotoxic agents, upregulation of the related placental growth factor (PlGF) presents a potential mechanism of resistance to VEGFA blockade in addition to being an independent risk factor for cancer progression [4–8]. Efforts to prevent all signaling through the VEGF receptors VEGFR1 and VEGFR2 using a soluble receptor-fusion protein with anti-PD-L1 resulted in increased immune filtration into pancreatic tumors but using receptor fusions are not without toxicity concerns [9,10]. Since PlGF is not required for normal angiogenesis, blockade of PlGF may have improved safety over existing treatments. However, the complexities of angiogenic signaling and the few publicly available antibodies targeting specific PlGF/receptor interactions have limited evaluation of this therapeutic strategy.

Tumor PlGF expression is independently associated with higher grade tumors, local recurrence, metastasis, and lower patient survival in a variety of cancers [11]. Furthermore, rectal and metastatic colorectal cancer patients exhibited elevated PlGF plasma levels following bevacizumab treatment [12,13], implicating PlGF as an angiogenic rescue pathway. Building on these observations, Fischer *et al*. reported an anti-PlGF antibody that prevented tumor growth and metastasis in multiple models [14], although a subsequent report challenged the generality of these results using additional antibodies and tumor models [15]. Mechanistic studies investigating the receptors and pathways responsible for anti-PlGF effects are inconclusive and contradictory [16,17]. Together, these reports highlight a need to define the impact of disrupting specific PlGF/receptor interactions to guide design of therapeutic strategies.

Antibodies are useful tools to disrupt specific protein-protein interactions and can provide mechanistic insights into the contributions of different receptors. PlGF binds the tyrosine kinase receptor VEGFR1 and the neuropilin co-receptors NRP1 and NRP2 but notably does not engage VEGFR2, which mediates canonical angiogenic VEGF effects. PlGF has been reported to exert pro-oncogenic and pro-metastatic activities through VEGFR1; PlGF also appears to mediate effects directly through NRP1, although this signaling cascade is less clear since NRPs do not contain a direct enzymatic intracellular signaling motif. PlGF, like VEGF, exists in several isoforms with differing receptor binding capabilities: the smaller isoforms comprise anti-parallel cysteine knot homodimers that bind VEGFR1 (*e*.*g*., VEGFA_121_ and human PlGF-1), while larger isoforms include a c-terminal heparin-binding domain that binds NRP receptors and is stabilized by heparin (*e*.*g*., VEGFA_165_, mouse and human PlGF-2; **Figure 1**).

**Figure 1.**
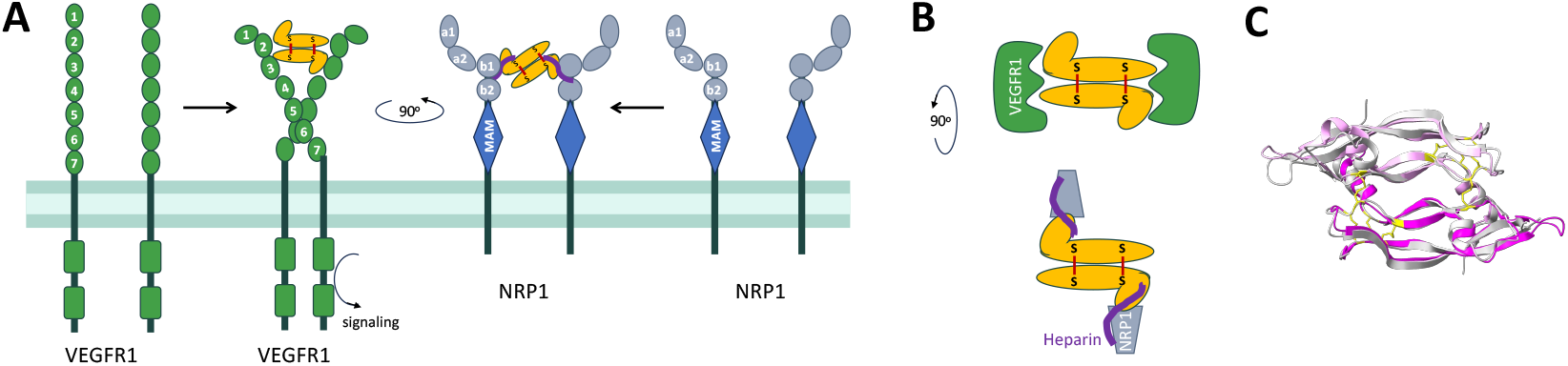
Binding of VEGFA_165_ and PlGF-2 to the VEGFR1 and NRP1 receptors. ***A***, Schematic of VEGF and PlGF interactions (yellow) with the intact VEGFR1 and NRP1 receptors. ***B***, A top-down view of the growth factor/ receptor complexes with heparin bridging the NRP1/ growth factor interface. **C**, Overlay of the core cysteine knot domains of VEGFA_121_ (grey, PDB 1RV6) and PlGF-1 (magenta, PDB 6T9D) to highlight structural similarities (RMSD 0.528 Å). Within the cysteine knot, loops L2 from protomer 1 and L1 and L3 from protomer 2 engage a single VEGFR1 or 2 receptor. The c-terminal heparin binding domains have only been resolved for VEGF_165_ and are not shown.

PlGF-binding antibodies that block interactions with VEGFR1 and NRP1 have been evaluated, including C9.V2 [15,16], 7A10 [15], 5D11D4 [14,18], and TB-403 [17], but their specific effects remain unclear and contradictory. Antibody C9.V2 primarily blocks PlGF binding to VEGFR1 and partially blocks NRP1 but only limits autocrine/ paracrine growth signals in cells expressing VEGFR1. Since C9.V2 treatment did not enhance survival in mouse xenograft models, the therapeutic relevance of its VEGFR1-dependent effects are unclear. Antibody TB-403 appears to potently inhibit the growth and spread of medulloblastoma in an NRP1-dependent manner *in vivo*, but data documenting its receptor blocking profile have not been published. Further complicating matters, while VEGFR1 and NRP1 are highly expressed on endothelial cells, they are both also expressed with varying cell surface densities on macrophages and tumor cells [19].

Antibodies specifically blocking individual PlGF/ receptor interactions could help to define the role of each receptor [15,17]. To address this need, we aimed to discover and characterize novel antibodies that bind PlGF to block its interactions with NRP1 in the presence of heparin. Anticipating that PlGF expression can be a resistance mechanism to anti-VEGF treatment, we also sought antibodies that target epitopes shared by PlGF and VEGFA to simultaneously inhibit both growth factors and limit resistance. This work identifies a panel of antibodies with distinct receptor blocking profiles to help define mechanisms of protection mediated by PlGF blockade.

## Materials & Methods

### Mouse Immunization

Three BALB/c mice (6 weeks old) were each immunized with 3 µgs of recombinant human PlGF-2 (PlGF-2, R&D Systems) and Freund’s complete adjuvant (Sigma Aldrich), then boosted 4 and 7 weeks later with another 3 µgs PlGF-2 and Freund’s incomplete adjuvant (Sigma Aldrich). Serum was collected at 0, 3, 4, 7, and 8 weeks to quantify the PlGF-2 antibody response and the mice were euthanized at 8 weeks and their spleens were collected in RNALater. All animal protocols were approved by the University of Texas at Austin Institutional Animal Care and Use Committee under IACUC protocol AUP-2018-00092 and mice were handled in accordance with IACUC guidelines.

### Library generation

Mouse spleens were homogenized and total RNA isolated using TRIzol (Invitrogen) and the PureLink RNA kit (Invitrogen) according to the manufacturers’ instructions. [20] First-strand cDNA synthesis was performed using 500 ng RNA and the SuperScript IV Transcriptase kit (Invitrogen) according to the manufacturer’s instructions and stored at –20 °C until further use. Degenerate primers were used to amplify the mouse antibody variable heavy (V_H_) and variable light (V_L_) chain domain-encoding fragments [21]; these were then combined into single-chain variable fragments (scFv) by overlap extension PCR and cloned into the pMopac24 pIII phage display vector [22] via directional *Sfi*I restriction sites for transformation into electrocompetent XL1-Blue *E. coli*.

### Phage display

M13K07 Helper Phage (New England Biolabs) were propagated in *E. coli* strain XL1-Blue, harvested, and used to infect *E. coli* containing the PlGF-2 scFv plasmid library. Phage were initially panned against an anti-myc tag antibody (ThermoFisher Scientific) to remove clones not expressing intact scFv-pIII fusion proteins. Next, the phage library was panned using several strategies: (1) screening against PlGF-1 (PeproTech) and repeated panning against PlGF-2 coats for six rounds, (2) alternative panning against PlGF-2 (R&D Systems) and mPlGF-2 (R&D Systems) coats for three rounds, and (3) alternative panning against PlGF-2 and VEGFA_165_ (R&D Systems) coats for 2-4 rounds. Recovered clones were screened as monoclonal phage in ELISAs to assess reactivity to wells coated with the anti-myc tag antibody 9E10, PlGF-2, VEGFA_165_ or uncoated wells.

### Protein expression constructs

Selected scFvs were sub-cloned for expression as full-length antibodies as previously described [23]. This included expression as chimeras comprising the mouse variable regions with human IgG1 and kappa chain constant regions, here called huFc. When a human Fc would interfere with the experiment, the heavy chain was altered such that the human C_H_1 domain was followed by the hinge, C_H_2 and C_H_3 domains from mouse IgG2a (called mFc) or the C_H_1 domain was appended with a stop codon to express human Fabs. The NRP1 receptor was obtained as a G-block and cloned into plasmid pcDNA3.1 (ThermoFisher Scientific) containing domains α1α2β1β2 fused in-frame with a c-terminal mFc as a purification handle, while the VEGFR1 domains 1-6 were cloned in-frame with a c-terminal huFc, both using AbVec antibody expression plasmids. [23] Growth factors human VEGFA_165_ and PlGF-2 were cloned into pcDNA3.1 plasmids with N-terminal Twin-Strep tags (IBA Lifesciences).

### Protein Expression and Purification

Antibodies and Fab fragments were transfected and expressed in ExpiCHO (ThermoFisher Scientific) for 7-10 days before harvest, while receptor-Fc fusion proteins were expressed in ExpiHEK (ThermoFisher Scientific) for 5 days before harvest, both according to the manufacturer’s instructions. Full length antibodies and receptor-Fc fusions were purified by an ÅKTA Pure FPLC system (Cytiva) via 1 mL HiTrap Protein A HP column (Cytiva) affinity chromatography. Fab proteins were purified by CaptureSelect CH1-XL Affinity Matrix (ThermoFisher Scientific) gravity chromatography. Growth factors expressing the Twin-Strep tag (IBA Lifesciences) were transfected and expressed in Freestyle 293F cells for 48 h before harvest. Growth factors were purified via Strep-Tactin XT resin (IBA Lifesciences) gravity chromatography at 4°C. In all cases, purified protein size, stability, and monodispersity were characterized by SDS-PAGE, Protein Thermal Shift Assay (ThermoFisher Scientific), and size exclusion chromatography with a Superdex 200 Increase 10/300GL column (Cytiva) on an ÅKTA Pure FPLC system.

### ELISA to assess antibody binding

Indirect enzyme linked immunosorbent assays (ELISAs) were performed using high-affinity 96-well plates (Corning) coated with antigen at 1 µg/mL and blocked with 5% milk in PBS with 0.05% Tween-20 (PBS-T). Coating antigens including commercial growth factors: PlGF-2, mPlGF-2, VEGFA_165_ (R&D Systems), and PlGF-1 (PeproTech). Purified antibodies were then serially diluted in 5% milk PBS-THuFc antibodies were detected with goat-anti-human Fc-HRP (Southern Biotech) at 1:1000 dilution in 50 µL. Plates were washed 3x after blocking, primary, and secondary incubation steps. ELISA signal was developed with 50 µL of TMB substrate (ThermoFisher Scientific) and quenched with 50 µL 1 N HCl after 1 min before reading the absorbance at 450 nm on a Molecular Devices SpectraMax M5 plate reader. Poly-specificity ELISAs were performed as described [24,25] using the following coat antigens: hemocyanin from *Megathura crentulata* (KLH) at 5 µg/mL (Sigma), single stranded DNA at 1 µg/mL (Sigma), double stranded DNA at 1 µg/mL (Sigma), insulin at 5 µg/mL (Sigma), each in PBS, and cardiolipin (Sigma) at 50 µg/mL in 100% ethanol.

Competition ELISAs were designed to assess antibody ability to bind the growth factor and block receptor binding, with and without the presence of heparin. Receptor-Fc fusion proteins, NRP1-huFc and VEGFR1-mFc, were biotinylated with an EZ-Link™ Sulfo-NHS-Biotinylation Kit (ThermoFisher Scientific) to provide a unique detection handle in this assay. ELISA plates were coated with PlGF-2 or VEGFA_165_ at 0.5 µg/mL and blocked with 3% bovine serum albumin in PBS with 0.05% Tween-20 for 1 h. Plates were washed 3x with PBS-T and then underwent primary incubation with biotinylated receptor (biotinylated VEGFR1-mFc at 1.6 ng/mL, biotinylated NRP1-mFc 250 ng/mL final concentration), competing antibody or non-biotinylated receptor at the indicated concentration and unfractionated heparin (ThermoFisher Scientific) for 1 h. Plates were washed 3x with PBS-T then biotinylated receptor detected with 1:1000 anti-streptavidin HRP (Sigma). Plates were washed a final 3x with PBS-T then developed with 50 µL TMB substrate for 20 min before quenching with 50 µL of 1 N HCl. High signal indicates that the competing protein did not impair growth factor binding to the receptor, while reduced signal indicates competition. Since the physiological concentration of heparin and heparan sulfates is highly variable based on tissue, we chose to use a high concentration of 1000 ng/mL that aligns with physiological levels in human plasma [26] and an intermediary concentration of 10 ng/mL, which is the concentration prescribed by Lonza’s endothelial growth medium to capture the relevant range but presume that 10 ng/mL heparin better represents extracellular matrix concentrations. [27,28] The inverted normalized competition ELISA signal at 10 µg/mL NRP1 or 20 µg/mL VEGFR1 are shown in Figure 2, where normalization was calculated using the following formula:

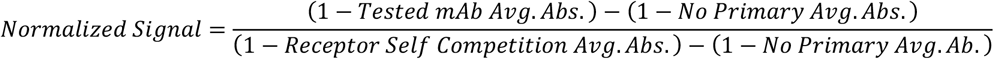

**Figure 2:**
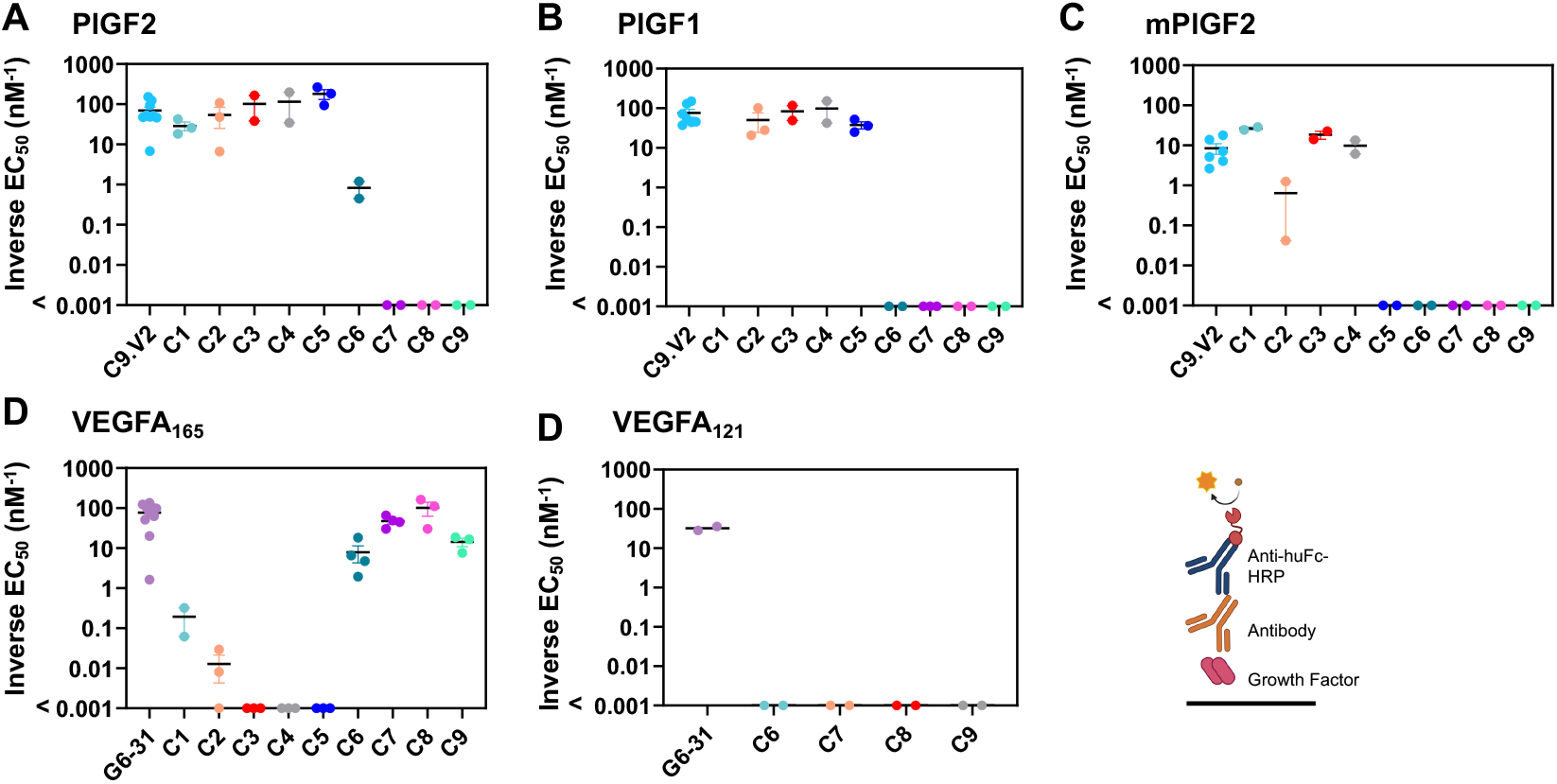
Selected antibodies bind PlGF and VEGFA isoforms. Antibodies C1-C9 were expressed as chimeric human IgG1/κ antibodies and evaluated for cytokine binding by ELISA against immobilized **A**, PlGF-2, **B**, PlGF-1 (which lacks the heparin-binding domain), and **C**, mPlGF-2 as well as **D**, VEGFA_165_ and **E**, VEGF_121_ (which lacks the heparin-binding domain) with inverted EC_50_ values reported. Data shown are the inverted EC_50_ from a four-parameter fit to a 7-point dilution curve, with bars reflecting the mean ± standard error; values listed as <0.001 nM were below the level of detection.

### Antibody binding kinetics & epitope binning

Kinetics and epitope binning were performed using biolayer interferometry (BLI, Forte Bio Octet). To measure equilibrium affinities, AHC tips (Sartorius) were used to capture anti-strep tag antibody C23.21-huFc at 200 nM, followed by incubation with Twin-Strep tagged growth factor at 200 nM, and then the IgG or Fab of interest at the concentration indicated. Antibodies were allowed to associate until the response curve stabilized, and Langmuir curves were generated by averaging the last ten datapoints per concentration measured and fitting them to a one-site specific binding model in GraphPad Prism to calculate a K_D_ value. All kinetic measurements were completed in duplicate. For epitope binning, AMC tips (Sartorius) were used to immobilize a capture antibody, mAb1-mFc, at 100 nM, followed by the growth factor (25 nM PlGF-2 or VEGFA; R&D Systems), and then exposed mAb1 site on the homodimer blocked by dipping into mAb1-huFc at 100 nM, before dipping the tips into a challenge antibody, mAb2-huFc at 100 nM. The measured signal shift was determined by subtracting the baseline response (nm) at 2310 s from that at 2810 s from step 3 mAb2-huFc association. In both studies, growth factor without final analyte was used as the subtracted baseline prior to analysis.

### In vitro cell culture

Human Umbilical Vein Endothelial Cells (HUVECs) constitutively expressing green fluorescent protein (GFP, Angio Proteomie) were grown in EGM-2 Complete Bullet Kit Media (Lonza) to 70% confluency; using only passages below 16. Four cancer lines (ATCC) were screened: Caki-I and H441 cell lines were cultured in 1640 RPMI, while MDA-MB-231 and MDA-MB-435S cell lines were cultured in DMEM, all supplemented with 10% FBS and 200 U/mL penicillin/streptomycin. All cells were grown in a humidified chamber at 37°C and 5% CO_2_. All cell lines were subject to monthly mycoplasma testing (Invitrogen) and were negative throughout the study period.

### HUVEC tube formation assay

96-well 3D µ-plates (Ibidi) that were filled with 10 µL of growth-factor-reduced Matrigel (Corning) on ice and then allowed to congeal at room temperature. HUVECs were then seeded at 5000 cells per well with antibody treatment of 10 – 0.01 µg/mL, with only 10 µg/mL used for the isotype. Antibodies were added to wells prior to cell seeding. Cells were incubated for 16-18 h, then imaged on an EVOS Cell Imaging System (ThermoFisher Scientific) via fluorescence and phase contrast. Tube formation was analyzed via ImageJ with the Angiogenesis Analyzer plugin [29]. The mesh number (formed rings) and branching interval (total segment length divided by the number of branched segments) were used to quantify tube formation. The study was repeated 3-4 times per antibody with 2-4 technical replicates each.

These data were standardized by checking for normality, subtracting the average value of the no antibody control and dividing by the standard deviation of the no antibody control for each measurement, per experiment.

### Flow cytometry to assess receptor display levels on cell lines

Cells were cultured until 70-80% confluency, washed once with PBS, then collected with a cell scraper (Fisher), and resuspended in blocking buffer (PBS with 10% FBS). Cells (3.5-5×10^5^) were incubated with a primary antibody at 15 ug/mL for 1 h and washed 3x with cold 10% FBS PBS and incubated with Alexa Fluor 647-labelled secondary antibodies (goat-anti-mouse-Fc-AF647 from Jackson ImmunoResearch or goat-anti-rabbit-Fc-AF647 from Invitrogen) at 1:1000 for 30 min and washed 3x with cold 10% FBS PBS before flow cytometry analysis using LSRFortessa SORP Flow Cytometer (BD) or an Attune NxT Flow Cytometer (ThermoFisher Scientific). Antibodies used include the monoclonal mouse anti-VEGFR2 antibody (R&D Systems), the polyclonal rabbit anti-VEGFR1 antibody (Invitrogen), the monoclonal mouse anti-NRP1 (Sino Biological) and an isotype control with mouse Fc (ITC-88 or M2B10). [30,31] Gates were set based on single cells with 1% overlap of Alexa Fluor 647 with the isotype control. Data were analyzed with FlowJo.

### Cell proliferation assay

Cells were passaged by scraping and then starved in serum-free DMEM (MDA-MB-231) or serum-free 1640 RPMI (Caki-I) for 24 h. The next day, cells were seeded at 5000 cells in 100 µL serum-deprived media per 96-well in tissue culture treated plates (TPP, CellTreat), with 6 technical replicates. While the cells settled, antibodies were pre-incubated with growth factors (PlGF-2 or VEGFA_165_, R&D Systems) for 1 h at 4°C. The antibody/ growth factor mixtures (100 µL) were added to wells, such that the final concentrations were 2.5 µg/mL or 5 µg/mL antibody and 12.5 ng/mL or 50 ng/mL growth factor. Cells were then allowed to grow for 72 h at 37°C with 5% CO_2_, at which point 20 µL of Resazurin (R&D Systems) was added to each well before an additional 48 h incubation. Fluorescence readings were collected on a SpectraMax M5 plate reader (Molecular Devices) at 555 nm excitation/ 585 nm emission or a Tecan Infinite 200 Pro M Plex Microplate reader at 560 nm excitation/ 590 nm emission. Data were normalized by dividing each result by the average of all control wells not treated with antibody or growth factor.

### Statistical analyses

All statistics were performed in GraphPad Prism. For the tube formation assay, first data was checked by Robust regression and OUTlier removal (ROUT) with a Q=1%. Then data were checked for normality by Shapiro-Wilk tests. If data was assumed to be normal and compared more than two groups, a one-way analysis of variance (ANOVA) followed by Dunnett’s multiple comparison test was completed. If not normal, a Kruskal-Wallis test followed by Dunn’s multiple comparison test was used.

### Data availability statement

The data generated in this study are available within the article and its supplementary data files and by request from the corresponding author.

## Results

### Discovery of antibodies binding human PlGF-2 and VEGFA_165_

In order to better define the role of the NRP1 receptor in mediating PlGF pathological effects, we identified a diverse set of high-affinity antibodies using immunized mice and phage display. We speculated that the PlGF receptor binding sites would be functionally constrained and less variable than the overall ∼60% sequence homology between PlGF and VEGF (**Supp. Fig. S1**). Accordingly, we hypothesized that PlGF-2 immunization could result in antibodies binding multiple PlGF and VEGF isoforms and designed selection strategies to identify antibodies targeting either growth factor. Since mouse models have indicated functional roles for human PlGF produced by the tumor xenograft as well as murine PlGF produced by stromal cells [2], we also aimed to identify antibodies exhibiting cross-species binding to human and mouse PlGF.

Balb/c mice were immunized with human PlGF-2, a splice-variant that includes the PlGF-1 isoform plus a 21-residue, c-terminal heparin- and NRP1-binding domain (**Supp Fig S1A**) [1], with antibody variable heavy and light chain regions amplified to generate an M13 p3-fusion library with 1.2×10^8^ unique members. First, to identify antibodies binding the NRP1-binding domain unique to PlGF-2, phage were panned against human PlGF-2, then screened to identify those lacking PlGF-1 binding. Second, to identify antibodies binding conserved regions that likely include receptor binding sites, phage were sequentially panned for binding to human and then mouse PlGF-2. Third, to select antibodies binding conserved epitopes within the NRP1- and VEGFR1-binding domains on both growth factors, phage were panned sequentially on human PlGF-2 and then human VEGFA_165_ **(Supp. Fig. S2**).

Clones recovered from each strategy were screened by monoclonal phage ELISA to assess binding to individual antigens (∼1000 total). This identified nine scFv clones with unique sequences and antigen binding profiles (C1-2 from strategy 1, C3-4 from strategy 2 and C5-C9 from strategy 3; **Supp Fig. S3**). These antibodies were all well-expressed as chimeric antibodies with human IgG1/ kappa constant domains (120-400 mg/L in ExpiCHO) and resulted in highly pure, monodisperse protein preparations, as assessed by size exclusion chromatography and SDS-PAGE (**Supp Fig. S3**). They also exhibited typical antibody thermal stabilities (≥68.8°C) with no observed binding to a panel of poly-specificity antigens [24,25]. In these experiments, the anti-PlGF control antibody C9.V2 consistently exhibited polyspecificity, characterized by binding to single stranded DNA, double stranded DNA and keyhole limpet hemocyanin (**Supp Fig. S3**). These nine chimeric antibodies were carried forward as lead candidates for further characterization.

### Identification of receptor-blocking antibodies

To compare antibody binding profiles, we performed ELISAs by immobilizing growth factor antigen followed by the addition of serially diluted antibodies to each well (**Fig. 2, Table 1, Supp Fig. S4**). As expected, the well-characterized anti-VEGF control antibody G6-31 bound its known ligands VEGFA_165_ and VEGFA_121_, while C9.V2 bound PlGF-1 and the human and mouse homologs of PlGF-2. Competition ELISAs were performed to evaluate the receptor-blocking profiles of each antibody by incubating biotinylated NRP1 or VEGFR1 in wells with immobilized growth factors in the presence or absence of each antibody. (**Fig. 3**). Antibody blockade was compared using two methods: (i) 50% inhibitory concentrations (IC_50_) and (ii) after normalizing one antibody concentration (10 mg/ml for NRP1, 20 mg/ml for VEGFR1) to receptor self-competition. For the second method, if no antibody was present, the response was 0, while receptor self-competition gave a response of 1.0. The control antibodies C9.V2 and G6-31 blocked receptor binding similarly as receptor self-competition using either metric (**Supp Fig S9**,**10**) [15,32]. Interestingly, although G6-31 acts as a VEGFR1 mimic by binding a structurally and energetically similar epitope on VEGFA [33], it also partially inhibited NRP1 recruitment. When low heparin concentrations were included, the G6-31 and C9.V2 antibodies less effectively blocked NRP1 binding, consistent with heparin stabilization of growth factor heparin-binding domain and its interaction with NRP1 [34].

**Table 1:** Antibody-antigen binding profiles*.

|  | G6-31 | C9.V2 | C1 | C2 | C3 | C4 | C5 | C6 | C7 | C8 | C9 |
| --- | --- | --- | --- | --- | --- | --- | --- | --- | --- | --- | --- |
| PIGF-2 | ND | +++ | +++ | +++ | +++ | +++ | +++ | + | - | - | - |
| mPIGF-2 | ND | +++ | +++ | + | +++ | +++ | - | - | - | - | - |
| PIGF-1 | ND | +++ | - | +++ | +++ | +++ | +++ | - | - | - | - |
| VEGFA<sub>165</sub> | +++ | ND | + | - | - | - | - | +++ | +++ | +++ | +++ |
| VEGFA<sub>121</sub> | +++ | ND | ND | ND | ND | ND | ND | - | - | - | - |

\*Relative antibody binding as measured by ELISA, with '+++’ indicating strong binding (EC<sub>50</sub> <100 μg/mL), ‘+’ indicating weak binding (signal did not saturate at 100 μg/mL), and ‘-’ indicating no observed binding; ND, not determined.

**Figure 3:**
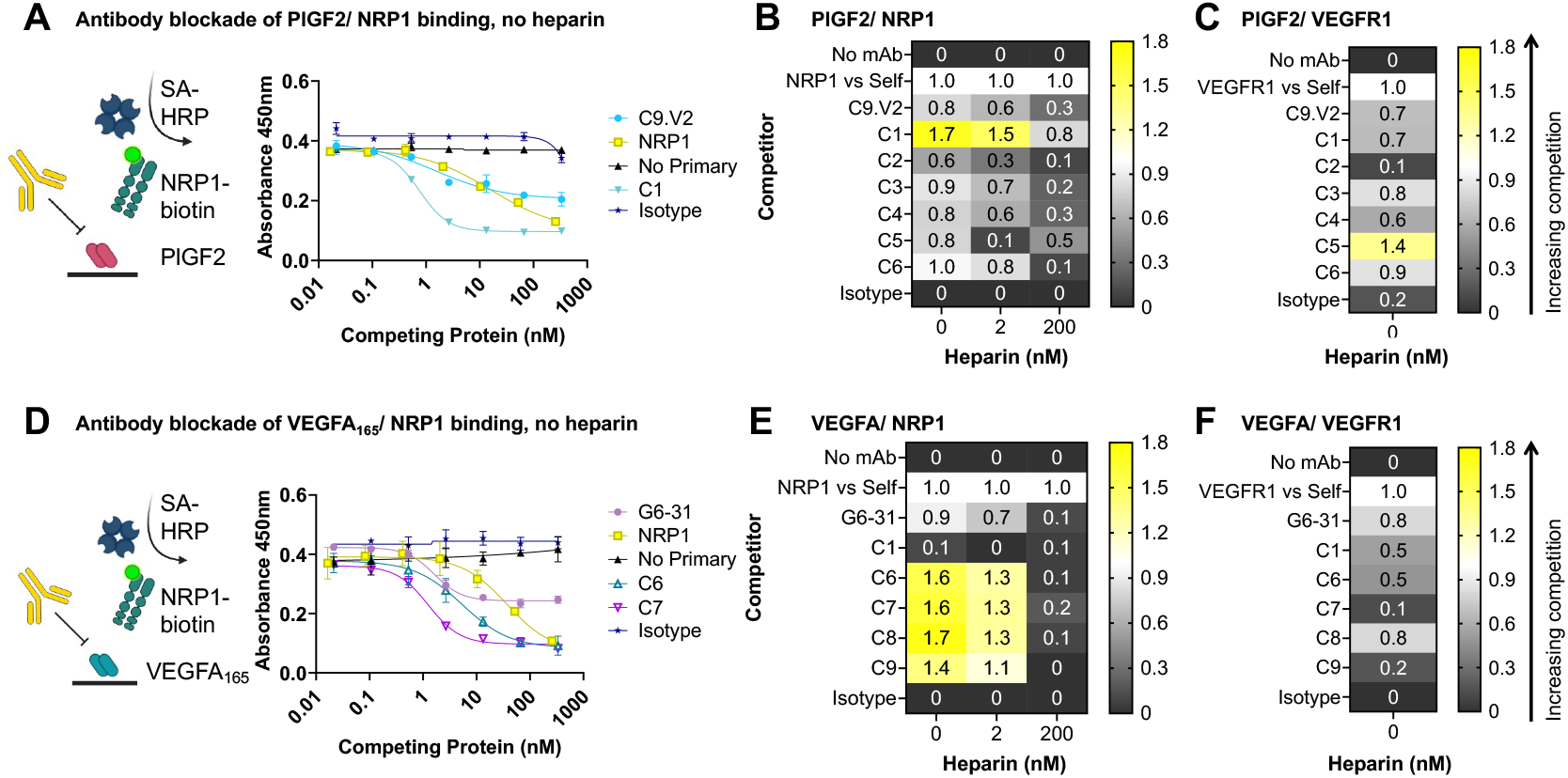
Cytokine/ receptor blocking profiles of selected antibodies. **A**, Sample competition ELISA data for antibody blockade of PlGF/ NRP1 interactions in the absence of heparin. Heat maps of the normalized competition ELISA signals for blockade of **B**, PlGF binding to 10 µg/mL NRP1 and **C**, PlGF binding to 20 µg/mL VEGFR1. **D**, Sample antibody competition ELISA data for blockade of VEGFA_165_/ NRP1 interactions in the absence of heparin. Heat maps of the normalized competition ELISA signals for blockade of **E**, VEGFA_165_ binding to 10 µg/mL NRP1 and **F**, VEGFA_165_ binding to 20 µg/mL VEGFR1.

Panning strategy 1 identified clone C1, which exhibited strong binding to mouse and human PlGF-2 but did not bind PlGF-1, and clone C2, which bound both PlGF isoforms (**Fig. 2)**. These profiles indicate that C1 binds residues in the exon 6 insert unique to PlGF-2, while C2 binds a shared epitope outside this region. Consistent with this, the receptor binding assay showed strong blockade of PlGF-2 binding to NRP1 by C1 (more potent than receptor self-blockade), while C2 had no effect (**Fig. 3)**. C1 inhibition of NRP1 binding was superior to C9.V2 even at high heparin concentrations (1000 μg/ml), while its impact on VEGFR1 binding was minimal (**Supp. Fig 10)**. Comparison of the NRP1 receptor blocking curves shows partial inhibition by C9.V2, suggesting it may block binding of just one of the two NRP1 receptor binding sites or exert an allosteric effect, while C1 fully inhibited NRP1 binding. Strategy 2 identified clones C3 and C4 with strong binding to both human PlGF isoforms and the mouse PlGF-2 homolog (**Fig. 2**). While C3 modestly blocked PlGF interactions with both receptors, NRP1 inhibition was diminished by high heparin concentrations, and it showed inferior blockade as compared to C1 (**Fig. 3**).

Strategy 3 used alternate panning against PlGF-2 and VEGFA_165,_ intended to identify antibodies binding conserved residues in the receptor binding sites. Despite immunizing with PlGF-2, clone C5 strongly bound both human PlGF isoforms, indicating it binds outside the NRP1 binding domain. Consistent with this, C5 blocked binding to VEGFR1 more potently than C9.V2 but showed a similar weak blockade of PlGF-2 binding to NRP1. By contrast, antibodies C6-C9 all bound VEGFA_165_ but not VEGFA_121_, indicating they bind epitopes in the NRP1/heparin-binding domain (**Fig. 2; Supp Fig S1**). These four antibodies strongly blocked VEGF_165_ binding to NRP1 in the presence of low heparin levels and minimally impacted VEGFR1 binding (**Fig. 3**). Only clone C6 exhibited dual-specificity, with modest binding to PlGF-2 and VEGFA_165_ and blocked both VEGF/ NRP1 and PlGF/ NRP1 interactions in the absence of heparin. This suggests that PlGF/ VEGF dual-specificity may come at the expense of high affinity to any single growth factor. Overall, we identified antibodies that selectively block PlGF/ NRP1 interactions (C1), PlGF/ VEGFR1 interactions (C5) and VEGFA/ NRP1 interactions (C6-C9), even in the presence of heparin.

To better understand how these antibodies bind their respective growth factors and the epitopes involved in receptor blockade, we used biolayer interferometry (BLI) to identify antibody competition groups. Antibodies were expressed with mouse Fc domains and captured on anti-mouse Fc BLI tips as mAb1-muFc and used to capture the growth factor. After blocking the second mAb1 site on the growth factor homodimer with mAb1-huFc, the ability to simultaneously bind a second antibody (mAb2-huFc), was measured. Modest BLI shifts of 0.7-0.9 nm were considered evidence of simultaneous binding due to partially overlapping epitopes, while larger shifts of >1.0 nm were considered evidence of simultaneous binding to distinct epitopes (**Fig. 4A**). When C9.V2 or C5 was used to capture PlGF-2, C1 yielded a strong signal, indicating C1 binds a different epitope. However, when C9.V2 was used for capture, C6 yielded a modest response, indicating these antibodies may bind overlapping epitopes. This dataset defined a potent NRP1-blocking site on PlGF-2 recognized by C1, a second, less potent NRP1-blocking epitope bound by C5 and C9.V2 and an overlapping sub-epitope recognized by C2-C4 that modestly inhibits both NRP1 and VEGFR1 binding (**Fig. 4E**). Antibodies G6-31 and C7 simultaneously bind VEGF, suggesting that G6-31 defines an epitope with modest VEGFR1 and NRP1 blocking behavior, while C7 defines a more potent and selective NRP1-blocking epitope. Antibody C7 blocks binding of C6, C8 and C9, indicating they all share similar epitopes involved in NRP1-binding (**Fig. 4F**). Overall, we defined two distinct epitopes on each growth factor: a potent NRP1-blocking epitope defined by our antibodies and a less potent NRP1-blocking epitope shared by our antibodies and previously described antibodies.

**Figure 4:**
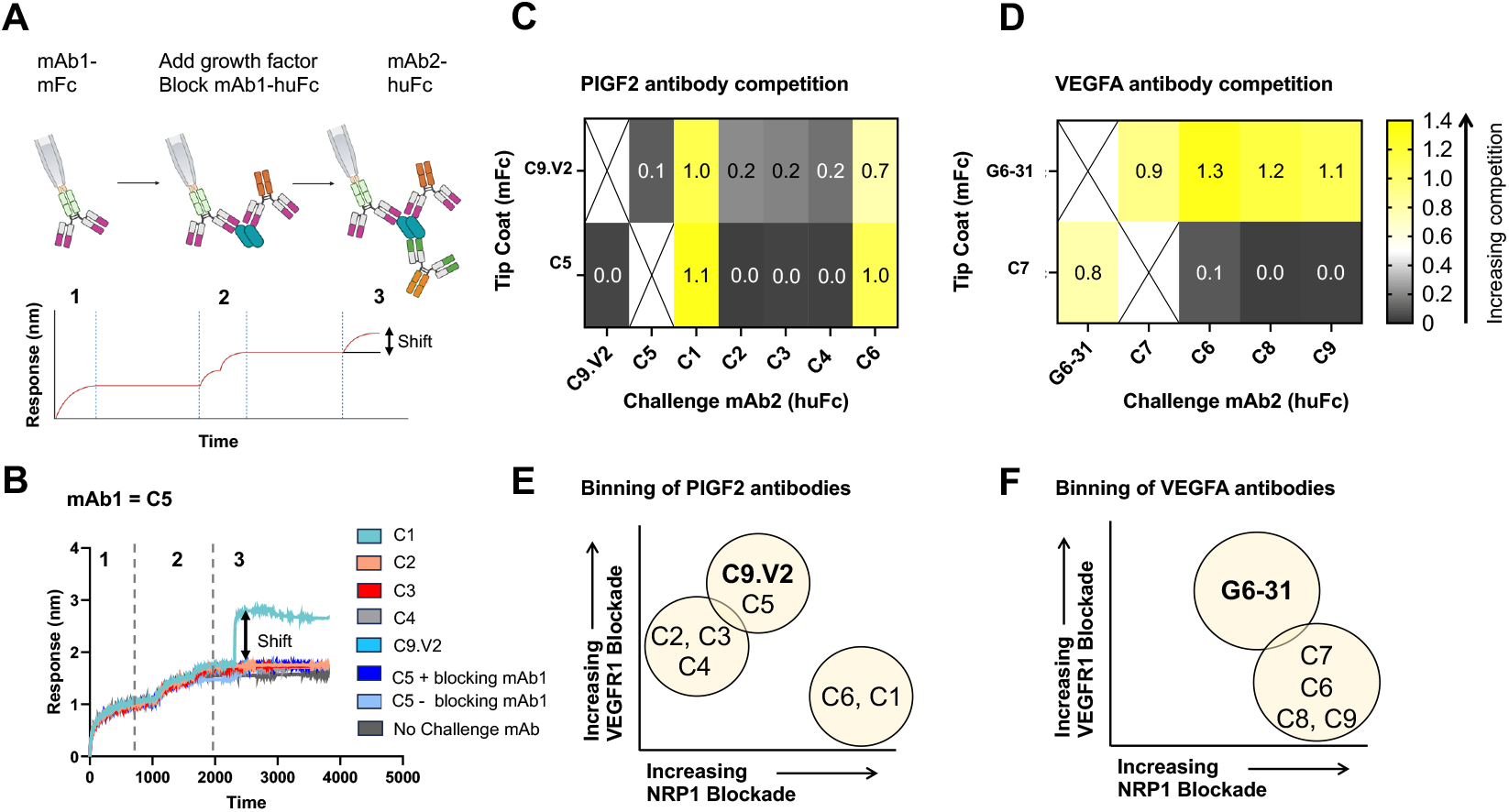
Competitive antibody binding assay reveals two separate epitopes per growth factor. **A**, Schematic of antibody binning strategy using BLI. **B**, Example BLI competition data showing the signal for mAb2 after 500 seconds association relative to the start of step 3 used to generate heat maps. Responses for antibodies binding to **C**, PlGF-2 after capture by C9.V2 or C5 and **D**, VEGFA_165_ after capture by G6-31 or C7. Venn diagrams summarizing antibody binning data for **E**, PlGF-2 and **F**, VEGFA_165_.

### Antibodies define potent receptor-blocking epitopes

Since antibody binding strength influences the ability to block receptor interactions and alter functional outcomes, we measured antibody equilibrium binding affinities (K_D_) using BLI. For this, we selected antibodies C9.V2 and G6-31 for benchmarking against prior reports, in addition to PlGF-binding antibodies C1, C2, and C5, and VEGF-binding antibodies C6 and C7 as representative antibodies from each unique epitope bin (**Table 2, Fig. S8-S11**). As bivalent antibodies, PlGF- and VEGF-binding antibodies all exhibited strong binding, with effective K_D_ values in the low nanomolar range. This corresponds well with prior reported K_D_ values of 0.8 nM for the C9.V2 IgG [15] and 0.9 nM and 0.2 nM for G6-31 Fab and IgG binding to VEGF_165,_ respectively, as measured by SPR. [32] Since several of these antibodies were observed to exhibit dual-reactive binding to PlGF-2 and VEGF_165_ by ELISA (**Fig. 1**), we also measured this using BLI. C1, which exhibited a 5.5 nM K_D_ for PlGF-2, showed negligible VEGF-binding at 200 nM, while C6 showed similar low nanomolar affinities for both growth factors (14 nM for PlGF-2 and 6.7 nM for VEGF_165_). Moreover, all antibodies except for G6-31 and C5 relied on bivalency for strong binding, as their measured Fab K_D_ values increased dramatically to ∼1000 nM. Since functional assays will use intact antibodies, the values measured for bivalent IgGs guided assay design even though their 2:2 binding stoichiometries will not reflect true K_D_ values.

**Table 2:** Antibody equilibrium K_D_ values measured by BLI*.

| *mAb* | *Binding to PIGF-2* |  |  |  |  | *Binding to VEGFA* |  |  |  |  |
| --- | --- | --- | --- | --- | --- | --- | --- | --- | --- | --- |
|  | C9.V2 | C1 | C2 | C5 | C6 | G6-31 | C1 | C2 | C6 | C7 |
| *Eff. $K_D$ (nM)* | 2.4<br>± 0.3 | 5.5<br>± 0.2 | 17.3<br>± 9.4 | 2.8<br>± 0.3 | 14.1<br>± 12.9 | 4.9<br>± 0.1 | NB | NB | 6.6<br>± 2.3 | 2.7<br>± 0.1 |
| *Avg. $R^2$* | 0.99 | 0.99 | 0.99 | 0.99 | 0.95 | 0.95 | NB | NB | 0.96 | 0.98 |
| ***Fab*** |  |  |  |  |  |  |  |  |  |  |
| *$K_D$ (nM)* | 728<br>± 67.8 | 1640<br>± 575 | 4680<br>± 2872 | 18.7<br>± 0.07 | NB | 47.6<br>± 19.1 | NB | NB | NB | 1038 ±<br>201 |
| *Avg. $R^2$* | 0.98 | 0.93 | 0.96 | 0.97 | NB | 0.95 | NB | NB | NB | 0.95 |

\*IgG values shown are effective $K_D$ (Eff. $K_D$ ), since antibodies and growth factors are dimers. $K_D$ value shown are average ± range, while NB indicates no binding.

### NRP1-blocking anti-VEGFA antibodies reduce angiogenesis

To assess antibody impact on VEGF-stimulated angiogenesis necessary for wound healing and other homeostatic functions, we used a HUVEC tube formation assay. Angiogenic tube formation assays are commonly used to assess vasculature regression exerted by VEGFA inhibitors [35,36], but PlGF inhibition in healthy tissues is thought to have a negligible effect. [37,38] As a result, we expected blockade of VEGFA/ NRP1 interactions to inhibit tube formation, while blockade of PlGF would have no effect. Cells were seeded on Matrigel overnight and imaged to enumerate tube formation by mesh number and branching interval to quantify angiogenesis (**Fig. 5**). [29] Cells grown in the presence of an isotype control antibody showed no difference versus untreated controls, whereas control antibody G6-31 which primarily blocks VEGFA-VEGFR1 and VEGFR2 interactions significantly inhibited both metrics at concentrations as low as 0.01 μg/mL. Antibody C6 inhibited mesh number under all doses but had minimal impact on branching interval, while C7 showed weak effects on both metrics at high concentrations (10 and 1 μg/ml). Notably, these antibodies bind distinct epitopes but share VEGF/NRP1-blocking activities (**Supp. Fig. S10)**. As expected, the anti-PlGF binding antibodies, including C9.V2, C1, C2 and C3 showed no effect at the highest antibody concentration tested (10 µg/mL). Overall, antibodies C6 and C7 support a role for NRP1 to contribute to VEGF-mediated angiogenesis.

**Figure 5:**
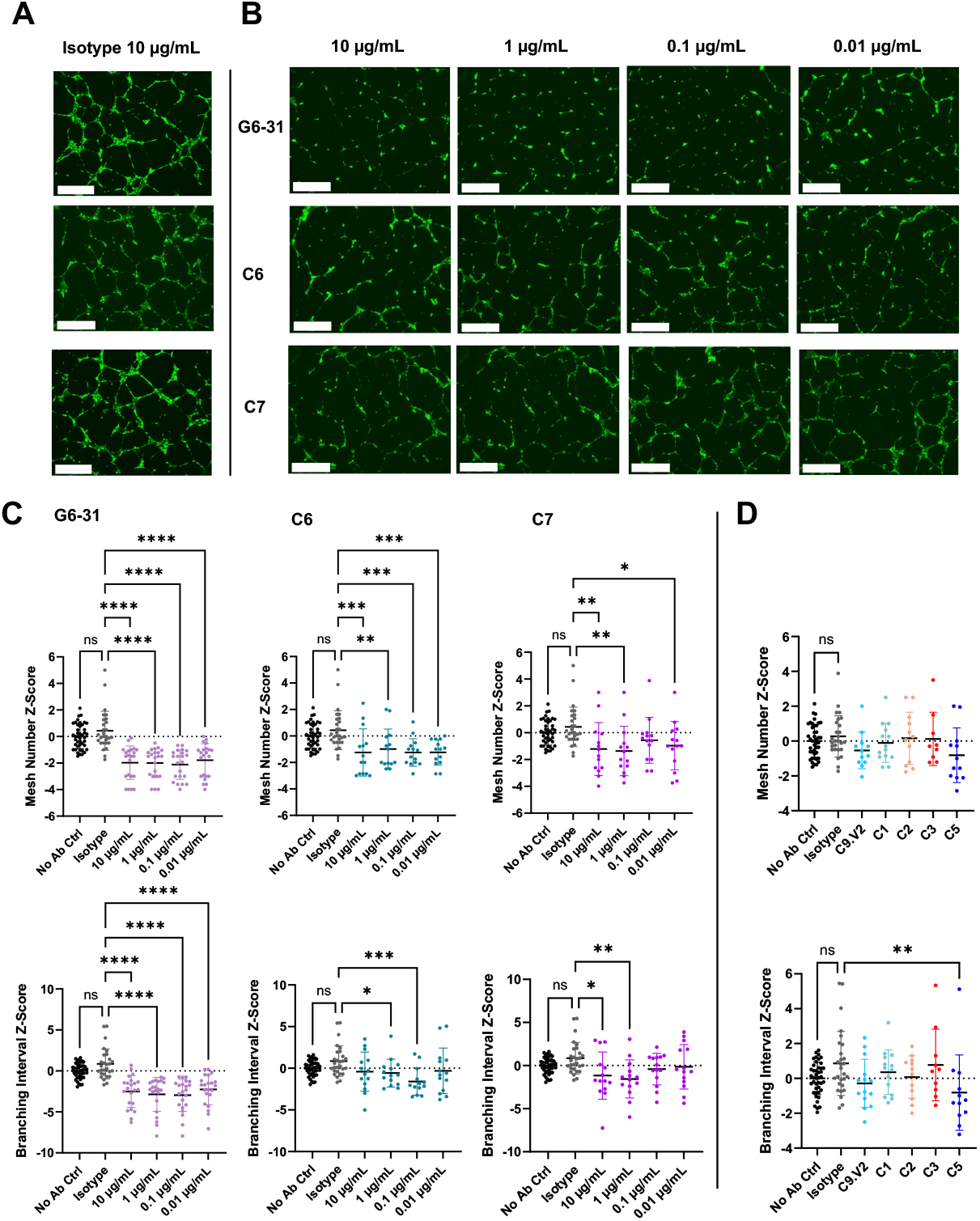
Potent NRP-1 blocking antibodies limit HUVEC tube formation. **A**, Qualitative GFP-fluorescent images of the isotype control dosed at 10 µg/mL for reference. Scale bar denotes 650 µm. **B**, Qualitative GFP-fluorescent images of antibody treated conditions with 10-fold dilutions starting at 10 µg/mL through 0.01 µg/mL. Conditions tested include the G6-31, C6, and C7; scale bar denotes 650 µm. **C**, Mesh number, also known as rings (top row), followed by branching interval, the total segment length divided by the number of branched segments (bottom row), standardized across experiments as a normal distribution Z-Score for each antibody conditions tested. **D**, Mesh number and branching interval measurements for PlGF antibodies at 10 µg/mL. Pairwise comparisons with the isotype control are indicated by * = p<0.05, ** = p<0.01, *** = p<0.001, and **** = p<0.0001).

### NRP1- and VEGFR1-blocking antibodies reduce Caki-I tumor cell proliferation

To assess antibody impact on pathological VEGFA and PlGF effects relevant for tumorigenesis, we developed a cell proliferation assay. Prior reports determined that expression of the relevant receptors is critical to detect PlGF-or VEGF-dependent tumor cell growth [16,17,39]. Accordingly, we screened four cancer cell lines (Caki-I [16], MDA-MB-231 [40,41], H441 [42,43], and MDA-MB-435S [44]) via flow cytometry to evaluate cell-surface receptor levels and assess susceptibility to these growth factors (**Fig S15A**). Caki-I and MDA-MB-231 cells showed low staining for VEGFR2 but high staining for VEGFR1 and NRP1 (**Fig. 6A, Supp. Fig. 20**), the two receptors of primary interest here, and were selected for use.

**Figure 6:**
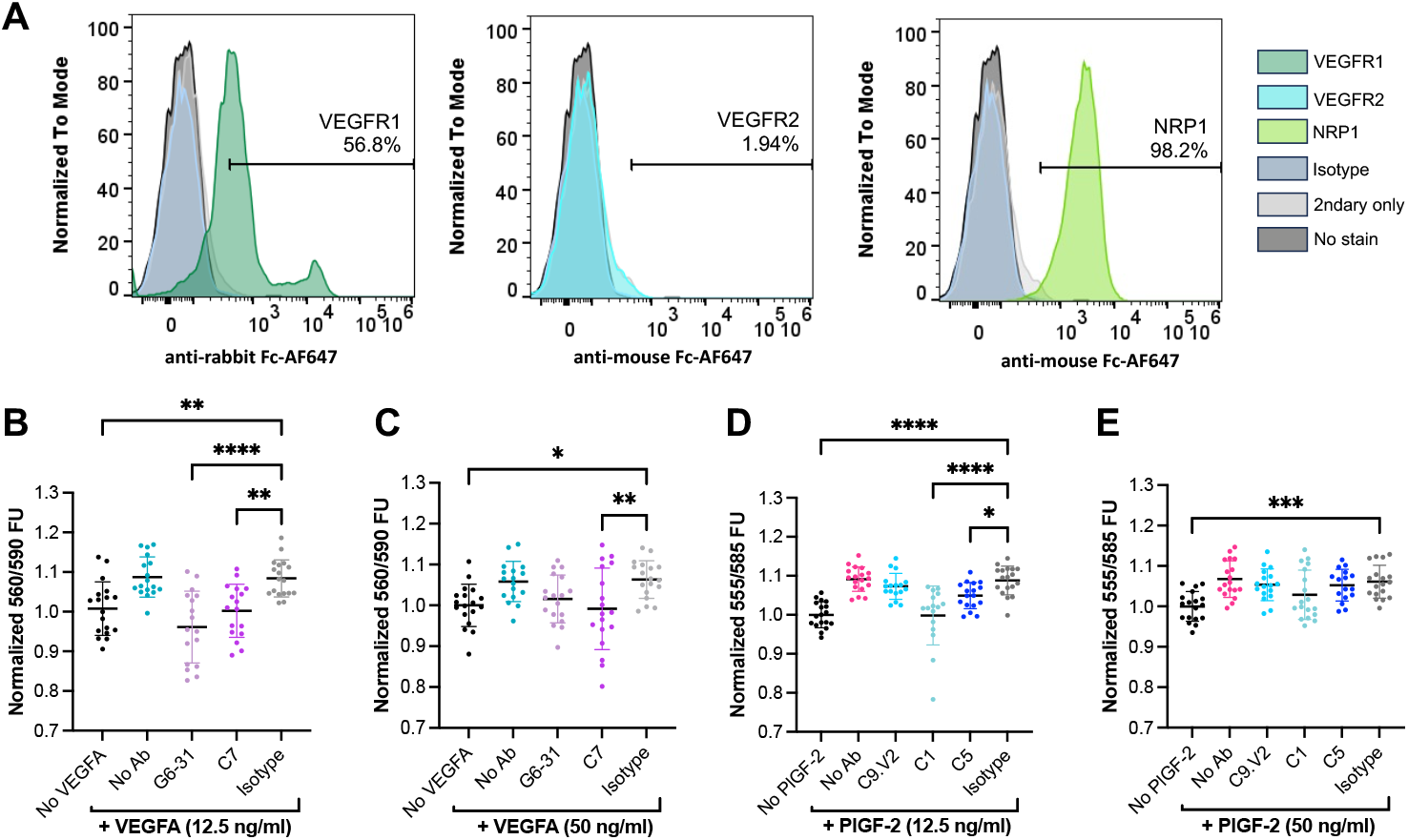
Antibodies blocking VEGFR1 and NRP1 interactions limit Caki-I cell proliferation. **A**, Caki-I clear cell carcinoma cells were screened for surface expression of VEGFR1, VEGFR2, and NRP1. Normalized Resazurin fluorescence attributed to Caki-I cells stimulated with VEGFA_165_ at **B**, 12.5 or **C**, 50 ng/mL, or PlGF-2 at **D**, 12.5 ng/mL or **E**, 50 ng/mL. In each case, the respective growth factor was pre-incubated with antibody (5 µg/mL for anti-VEGF, 2.5 µg/mL for anti-PlGF). Pairwise comparisons from ANOVA are indicated by * = p<0.05, ** = p<0.01, *** = p<0.001, and **** = p<0.0001).

These cells were evaluated in a proliferation assay to measure growth factor-mediated metabolic stimulation after serum starvation to enhance responses. This assay was based on an industrial assay to evaluate biological PlGF-2 activity with MDA-MB-231 cells in low-serum conditions [45]; however, the MDA-MD-231 cells were not strongly stimulated by PlGF-2 or VEGFA_165_ concentrations up to 50 ng/mL (**Fig S12B)**. By contrast, Caki-I cells demonstrated significant growth increase in the presence of PlGF-2 (∼7-9%) or VEGF (∼6-8%). When antibodies were pre-incubated with the growth factor, the isotype control and C9.V2 antibodies showed no impact on PlGF-2-stimulated growth, while C1 completely inhibited and C5 modestly inhibited growth (**Fig 4B,C**). These antibodies have similar binding affinities for PlGF-2 (5.5 and 2.8 nM effective K_D_, respectively; **Table 2**) but C1 primarily blocks PlGF binding to NRP1, while C5 primarily blocks PlGF binding to VEGFR1 (**Fig. 3**). These data indicate that NRP1 contributes to PlGF-2 activities in this assay. Similarly, the isotype control antibody showed no impact on VEGF-mediated growth, while G6-31 showed a modest impact and C7 strongly inhibited growth (**Fig 6D,E**). Since these antibodies also have similar affinities (4.9 and 2.7 nM K_D_, respectively; **Table 1**), but C7 strongly blocks NRP1, we conclude that NRP1 contributes to VEGF-dependent growth in this assay. Overall, our results support a role for NRP1 in mediating tumor cell growth stimulated by both PlGF and VEGF.

## Discussion

The clinical performance of anti-VEGF therapeutics has motivated interest in understanding the contributions of PlGF to pathological angiogenesis and the roles of their shared receptors in mediating biological activities. Similarly, recent observations of synergies between anti-PD-1/PD-L1 and anti-VEGF provides opportunities to include novel antibodies with tailored mechanisms of action into future treatments [10,46]. Here, we report the discovery of three antibodies disrupting these interactions: C1, which mediates highly selective blockade of PlGF-2/ NRP1 interactions in the presence of heparin to limit Caki-I cell proliferation; C7 which binds the heparin-binding domain on VEGFA_165_ to block NRP1 interactions, reduce HUVEC tube formation and Caki-I cell proliferation; and C5 which potently blocks PlGF/ VEGFR1 interactions and modestly blocks PlGF-2/ NRP1 interactions to exert a weak effect on Caki-I cell proliferation. In addition, we identified one clone with dual VEGFA_165_/PlGF-2 specificity (C6) that modestly inhibits VEGFA_165_/ NRP1 interactions, suggesting these conserved epitopes are harder to target, perhaps because they are composed of a small number of shared residues. These antibodies were discovered using distinct selection strategies, define novel binding sites on their growth factor targets, and support a role for NRP1 in mediating VEGFA_165_ and PlGF-2 activities.

A series of interactions and conformational changes between the growth factors and their receptors and ligands ultimately result in the signaling required to mediate PlGF and VEGF effects. This process offers multiple opportunities for antibody intervention (**Fig. 1**) and the antibodies reported here define three distinct mechanisms. First, the shared VEGFR1/2 footprint on VEGFA is well-defined, with bevacizumab and G6-31 both acting as receptor mimics to sterically block this interaction. Thus, it is not surprising that we isolated antibody C5, which blocks PlGF activities using a similar mechanism: C5 binds an epitope overlapping with C9.V2 on PlGF-1 and PlGF-2, suggesting it binds the core cysteine knot domain, likely near the dimer groove, to block VEGFR1 interactions. Both antibodies also partially block NRP1 binding, perhaps by adopting an angle of approach that prevents simultaneous binding to two NRP1 receptors or indirectly disrupts complex formation through allosteric effects (**Fig 7**). Notably, C5 more potently blocks VEGFR1 binding and PlGF activities than C9.V2 (**Fig. 7D**), which may reflect its stronger monovalent Fab affinity (18.7 nM K_D_ for C5, versus 728 nM for C9.V2) to PlGF-2. Second, NRP1 binds to the C-terminal domains of the growth factors independent of VEGFR1, an interaction that is strongly influenced by heparin. This negatively-charged glucosamino-glycan binds positively charged surface patches on PlGF-2 and VEGFA_165_ to neutralize electrostatic repulsion; heparin may also reduce the intrinsic disorder of this domain to support NRP1 binding [47]. Antibody C1 binds PlGF-2 to potently block NRP1 binding and inhibit Caki-I cell proliferation. Since C1 has highly charged heavy chain CDRs and its effects are only partially impaired by high heparin concentrations (1000 μg/ml), we speculate it binds near the heparin binding site. Since heparin does not block C1 binding to PlGF-2, C1 may stabilize an alternate confirmation or otherwise occlude receptor binding residues near the PlGF-2 C-terminus (**Fig. 8B**). Third, antibodies C6-C9 are potent inhibitors of VEGFA_165_/ NRP1 binding that map to a shared epitope on the C-terminal domain. C7 showed comparable biological activities as the VEGFR1-mimic antibody G6-31, despite their divergent binding sites and receptor-blockade profiles. Unlike C1, these antibody CDRs are less charged and receptor blocking is inhibited by high heparin concentrations. Accordingly, they may bind sites partially overlapping with the heparin binding sites on VEGFA_165_ to also occlude access to the peptide terminus that directly interacts with NRP1 (**Fig. 8C**). Overall, we identified antibodies blocking receptor binding by multiple mechanisms including direct steric blockade of a single receptor, indirect disruption of complex formation, and possible allosteric effects to impair conformational changes required for full signaling.

**Figure 7.**
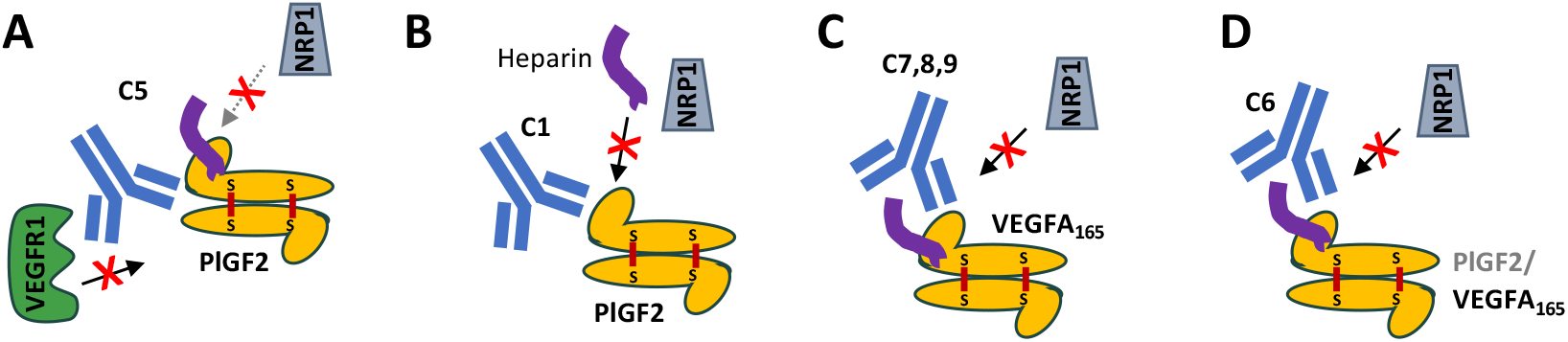
Schematics depicting suggested antibody mechanisms of activity. **A**, Antibody C5 binds a PlGF-2 epitope that potently blocks VEGFR1 binding, even as a monovalent Fab, to mediate weak biological activities. **B**, Antibody C1 acts potently prevents NRP1 binding to PlGF-2. **C**, Antibodies C7, 8 and 9 bind similar epitopes on the C-terminal domain of VEGFA165 to block NRP1 binding. **D**, Antibody C6 exhibits dual-reactivity by binding the heparin binding domain to block NRP1 interactions.

Due to their overlapping biological roles, PlGF-2 expression presents a potential resistance pathway in response to treatment with anti-VEGF antibodies. However, the shared structures and ligands (VEGFR1, NRP1, NRP2 and heparin) present opportunities to identify antibodies with dual-specificity to block both growth factors and thereby limit a mechanism of tumor resistance. NRP1 ligands follow a *C-end R* rule [48], which requires an extreme C-terminal arginine residue on ligands; accordingly, VEGFA_165_ and PlGF-2 share similar C-terminal sequences (CDKRPRR and AVPRR, respectively; **Supp Fig. S1**). Since C6 exhibits dual-reactive binding to VEGFA_165_ and PlGF-2 and blocks NRP1 interactions, we suspect this antibody binds conserved residues near the terminal peptide to alter its presentation or sterically inhibit receptor binding. Indeed, AlphaFold3 confidently predicts the docking of C6 Fab to the PlGF-2 and VEGFA_165_ heparin binding domains, with predicted contacts between V_H_ and the C-terminal sequences plus additional upstream contacts between both antibody chains and α-helices on each growth factor (**Fig. 8D, Supporting Fig 21**). The heparin binding sites present another potential target for dual-reactive antibodies, as these highly charged basic patches include conserved BBXB heparin-binding motifs, where B is a basic residue and X is any residue. An antibody binding near these sites could directly block heparin binding to prevent heparin-mediated stabilization of NRP1 interactions. Supporting this concept, antibody C1, which appears to act as a heparin mimic to bind PlGF-2, also exhibits weak VEGF binding likely due to charge-charge interactions from heparin-binding sites on both growth factors. Finally, while we did not discover any antibody with this behavior, several key hot spot residues mediating interactions with VEGFR1 are conserved across VEGFA_165_ and PlGF-2 (F17, E64, Q79, I83) and may form a shared epitope for future antibody discovery efforts. Similar strategies have been used to identify an antibody binding all members of the *Staphylococcal aureus* leukotoxin family which share even lower sequence homology (∼25%), although CDR engineering of an initial, partially multi-reactive hit was required. [49] Accordingly, further engineering of C1 or C6 may strengthen CDR interactions with the shared epitope to increase dual-specificity and result in a single antibody able to block NRP1 binding of each growth factor.

To better define the role of the NRP1 receptor in angiogenesis, we report the discovery of antibodies C1 and C7 which bind PlGF-2 and VEGFA_165_, respectively, in their respective NRP1-binding domains. These antibodies each demonstrated potent biochemical inhibition of receptor binding and biological activity in at least one *in vitro* activity assay. Furthermore, antibody C6 supports the potential for dual-reactive antibodies able to bind VEGFA_165_ and PlGF-2 by targeting homology within their NRP1-binding domains. The absence of structural data for either growth factor in the presence of heparin and lack of well-developed *in vitro* assays to evaluate PlGF-2 activities limited our ability to define antibody mechanisms of action. Precise identification of the antibody binding sites and their effects on receptor complex formation will contribute to the utility of these antibodies. Regardless, these antibodies add to a growing arsenal that can help dissect mechanisms of anti-VEGFA evasion, clarify the complex interplay between both growth factors, and support development of effective strategies that prevent pro-angiogenic signals and promote tumor vascular normalization – features that may contribute to design of future multi-functional antibodies that include anti-VEGF/ anti-PlGF components.

## Acknowledgements

This work was funded by the Cancer Prevention and Research Institute of Texas (RP190653 and RP220587 to J.A.M.), the Welch Foundation (F-1767 to J.A.M.) and NSF GRFP fellowhips to A.N.Q and S.A.B. We acknowledge the use of the CPRIT Advanced Protein Therapeutics core, RRID SCR_023740 for assistance with antibody production and characterization.

## Conflicts of Interest

S.A.B, A.N.Q., N.V.J. and A.W.N., J.A.M. are inventors on a patent disclosure for antibodies C1, C5, C6, C7, C8.

## References

[1] D.F. McDermott, M.A. Huseni, M.B. Atkins, R.J. Motzer, B.I. Rini, B. Escudier, L. Fong, R.W. Joseph, S.K. Pal, J.A. Reeves, M. Sznol, J. Hainsworth, W.K. Rathmell, W.M. Stadler, T. Hutson, M.E. Gore, A. Ravaud, S. Bracarda, C. Suárez, R. Danielli, V. Gruenwald, T.K. Choueiri, D. Nickles, S. Jhunjhunwala, E. Piault-Louis, A. Thobhani, J. Qiu, D.S. Chen, P.S. Hegde, C. Schiff, G.D. Fine, T. Powles, Clinical activity and molecular correlates of response to atezolizumab alone or in combination with bevacizumab versus sunitinib in renal cell carcinoma, Nat. Med. 24 (2018) 749–757. 10.1038/s41591-018-0053-3.

[2] F.S. Hodi, D. Lawrence, C. Lezcano, X. Wu, J. Zhou, T. Sasada, W. Zeng, A. Giobbie-Hurder, M.B. Atkins, N. Ibrahim, P. Friedlander, K.T. Flaherty, G.F. Murphy, S. Rodig, E.F. Velazquez, M.C. Mihm, S. Russell, P.J. DiPiro, J.T. Yap, N. Ramaiya, A.D. Van den Abbeele, M. Gargano, D. McDermott, Bevacizumab plus ipilimumab in patients with metastatic melanoma, Cancer Immunol. Res. 2 (2014) 632–642. 10.1158/2326-6066.CIR-14-0053.

[3] R.S. Finn, S. Qin, M. Ikeda, P.R. Galle, M. Ducreux, T.-Y. Kim, M. Kudo, V. Breder, P. Merle, A.O. Kaseb, D. Li, W. Verret, D.-Z. Xu, S. Hernandez, J. Liu, C. Huang, S. Mulla, Y. Wang, H.Y. Lim, A.X. Zhu, A.-L. Cheng, Atezolizumab plus Bevacizumab in Unresectable Hepatocellular Carcinoma, N. Engl. J. Med. 382 (2020) 1894–1905. 10.1056/NEJMoa1915745.

[4] Y. Yang, Y. Cao, The impact of VEGF on cancer metastasis and systemic disease, Semin. Cancer Biol. 86 (2022) 251–261. 10.1016/j.semcancer.2022.03.011.

[5] Y. Yang, Y. Zhang, H. Iwamoto, K. Hosaka, T. Seki, P. Andersson, S. Lim, C. Fischer, M. Nakamura, M. Abe, R. Cao, P.V. Skov, F. Chen, X. Chen, Y. Lu, G. Nie, Y. Cao, Discontinuation of anti-VEGF cancer therapy promotes metastasis through a liver revascularization mechanism, Nat. Commun. 7 (2016) 12680. 10.1038/ncomms12680.

[6] Y. Xu, Q. Li, X.-Y. Li, Q.-Y. Yang, W.-W. Xu, G.-L. Liu, Short-term anti-vascular endothelial growth factor treatment elicits vasculogenic mimicry formation of tumors to accelerate metastasis, J. Exp. Clin. Cancer Res. CR 31 (2012) 16. 10.1186/1756-9966-31-16.

[7] G.W. Prager, M. Poettler, M. Unseld, C.C. Zielinski, Angiogenesis in cancer: Anti-VEGF escape mechanisms, Transl. Lung Cancer Res. 1 (2012) 14–25. 10.3978/j.issn.2218-6751.2011.11.02.

[8] C. Montemagno, G. Pagès, Resistance to Anti-angiogenic Therapies: A Mechanism Depending on the Time of Exposure to the Drugs, Front. Cell Dev. Biol. 8 (2020) 584. 10.3389/fcell.2020.00584.

[9] P. Rougier, H. Riess, R. Manges, P. Karasek, Y. Humblet, C. Barone, A. Santoro, S. Assadourian, L. Hatteville, P.A. Philip, Randomised, placebo-controlled, double-blind, parallel-group phase III study evaluating aflibercept in patients receiving first-line treatment with gemcitabine for metastatic pancreatic cancer, Eur. J. Cancer Oxf. Engl. 1990 49 (2013) 2633–2642. 10.1016/j.ejca.2013.04.002.

[10] D.K. Kim, J. Jeong, D.S. Lee, D.Y. Hyeon, G.W. Park, S. Jeon, K.B. Lee, J.-Y. Jang, D. Hwang, H.M. Kim, K. Jung, PD-L1-directed PlGF/VEGF blockade synergizes with chemotherapy by targeting CD141+ cancer-associated fibroblasts in pancreatic cancer, Nat. Commun. 13 (2022) 6292. 10.1038/s41467-022-33991-6.

[11] M. Dewerchin, P. Carmeliet, Placental growth factor in cancer, Expert Opin. Ther. Targets 18 (2014) 1339–1354. 10.1517/14728222.2014.948420.

[12] E. Van Cutsem, C. Paccard, M. Chiron, J. Tabernero, Impact of Prior Bevacizumab Treatment on VEGF-A and PlGF Levels and Outcome Following Second-Line Aflibercept Treatment: Biomarker Post Hoc Analysis of the VELOUR Trial, Clin. Cancer Res. 26 (2020) 717–725. 10.1158/1078-0432.CCR-19-1985.

[13] R.K. Jain, D.G. Duda, J.W. Clark, J.S. Loeffler, Lessons from phase III clinical trials on anti-VEGF therapy for cancer, Nat. Clin. Pract. Oncol. 3 (2006) 24–40. 10.1038/ncponc0403.

[14] C. Fischer, B. Jonckx, M. Mazzone, S. Zacchigna, S. Loges, L. Pattarini, E. Chorianopoulos, L. Liesenborghs, M. Koch, M. De Mol, M. Autiero, S. Wyns, S. Plaisance, L. Moons, N. van Rooijen, M. Giacca, J.-M. Stassen, M. Dewerchin, D. Collen, P. Carmeliet, Anti-PlGF Inhibits Growth of VEGF(R)-Inhibitor-Resistant Tumors without Affecting Healthy Vessels, Cell 131 (2007) 463–475. 10.1016/j.cell.2007.08.038.

[15] C. Bais, X. Wu, J. Yao, S. Yang, Y. Crawford, K. McCutcheon, C. Tan, G. Kolumam, J.-M. Vernes, J. Eastham-Anderson, P. Haughney, M. Kowanetz, T. Hagenbeek, I. Kasman, H.B. Reslan, J. Ross, N. Van Bruggen, R.A.D. Carano, Y.-J.G. Meng, J.-A. Hongo, J.-P. Stephan, M. Shibuya, N. Ferrara, PlGF Blockade Does Not Inhibit Angiogenesis during Primary Tumor Growth, Cell 141 (2010) 166–177. 10.1016/j.cell.2010.01.033.

[16] J. Yao, X. Wu, G. Zhuang, I.M. Kasman, T. Vogt, V. Phan, M. Shibuya, N. Ferrara, C. Bais, Expression of a functional VEGFR-1 in tumor cells is a major determinant of anti-PlGF antibodies efficacy, Proc. Natl. Acad. Sci. U. S. A. 108 (2011) 11590–11595. 10.1073/pnas.1109029108.

[17] M. Snuderl, A. Batista, N.D. Kirkpatrick, C.R. de Almodovar, L. Riedemann, E.C. Walsh, R. Anolik, Y. Huang, J.D. Martin, W. Kamoun, E. Knevels, T. Schmidt, C.T. Farrar, B.J. Vakoc, N. Mohan, E. Chung, S. Roberge, T. Peterson, C. Bais, B.H. Zhelyazkova, S. Yip, M. Hasselblatt, C. Rossig, E. Niemeyer, N. Ferrara, M. Klagsbrun, D.G. Duda, D. Fukumura, L. Xu, P. Carmeliet, R.K. Jain, Targeting placental growth factor/neuropilin 1 pathway inhibits growth and spread of medulloblastoma, Cell 152 (2013) 1065–1076. 10.1016/j.cell.2013.01.036.

[18] S. Van de Veire, I. Stalmans, F. Heindryckx, H. Oura, A. Tijeras-Raballand, T. Schmidt, S. Loges, I. Albrecht, B. Jonckx, S. Vinckier, C. Van Steenkiste, S. Tugues, C. Rolny, M. De Mol, D. Dettori, P. Hainaud, L. Coenegrachts, J.-O. Contreres, T. Van Bergen, H. Cuervo, W.-H. Xiao, C. Le Henaff, I. Buysschaert, B.K. Masouleh, A. Geerts, T. Schomber, P. Bonnin, V. Lambert, J. Haustraete, S. Zacchigna, J.-M. Rakic, W. Jiménez, A. Noël, M. Giacca, I. Colle, J.-M. Foidart, G. Tobelem, M. Morales-Ruiz, J. Vilar, P. Maxwell, S.A. Vinores, G. Carmeliet, M. Dewerchin, L. Claesson-Welsh, E. Dupuy, H. Van Vlierberghe, G. Christofori, M. Mazzone, M. Detmar, D. Collen, P. Carmeliet, Further Pharmacological and Genetic Evidence for the Efficacy of PlGF Inhibition in Cancer and Eye Disease, Cell 141 (2010) 178–190. 10.1016/j.cell.2010.02.039.

[19] M. Tyler, A. Gavish, C. Barbolin, R. Tschernichovsky, R. Hoefflin, M. Mints, S.V. Puram, I. Tirosh, The Curated Cancer Cell Atlas provides a comprehensive characterization of tumors at single-cell resolution, Nat. Cancer 6 (2025) 1088–1101. 10.1038/s43018-025-00957-8.

[20] A.C. Patterson-Orazem, A.N. Qerqez, L.R. Azouz, M.T. Ma, S.E. Hill, Y. Ku, L.A. Schildmeyer, J.A. Maynard, R.L. Lieberman, Recombinant antibodies recognize conformation-dependent epitopes of the leucine zipper of misfolding-prone myocilin, J. Biol. Chem. 297 (2021) 101067. 10.1016/j.jbc.2021.101067.

[21] A. Krebber, S. Bornhauser, J. Burmester, A. Honegger, J. Willuda, H.R. Bosshard, A. Plückthun, Reliable cloning of functional antibody variable domains from hybridomas and spleen cell repertoires employing a reengineered phage display system, J. Immunol. Methods 201 (1997) 35–55. 10.1016/S0022-1759(96)00208-6.

[22] A. Hayhurst, S. Happe, R. Mabry, Z. Koch, B.L. Iverson, G. Georgiou, Isolation and expression of recombinant antibody fragments to the biological warfare pathogen Brucella melitensis, J. Immunol. Methods 276 (2003) 185–196. 10.1016/s0022-1759(03)00100-5.

[23] A.W. Nguyen, E.K. Wagner, J.R. Laber, L.L. Goodfield, W.E. Smallridge, E.T. Harvill, J.F. Papin, R.F. Wolf, E.A. Padlan, A. Bristol, M. Kaleko, J.A. Maynard, A cocktail of humanized anti-pertussis toxin antibodies limits disease in murine and baboon models of whooping cough, Sci. Transl. Med. 7 (2015) 316ra195. 10.1126/scitranslmed.aad0966.

[24] T. Jain, T. Sun, S. Durand, A. Hall, N.R. Houston, J.H. Nett, B. Sharkey, B. Bobrowicz, I. Caffry, Y. Yu, Y. Cao, H. Lynaugh, M. Brown, H. Baruah, L.T. Gray, E.M. Krauland, Y. Xu, M. Vásquez, K.D. Wittrup, Biophysical properties of the clinical-stage antibody landscape, Proc. Natl. Acad. Sci. U. S. A. 114 (2017) 944–949. 10.1073/pnas.1616408114.

[25] S.S. Pierangeli, E.N. Harris, A protocol for determination of anticardiolipin antibodies by ELISA, Nat. Protoc. 3 (2008) 840–848. 10.1038/nprot.2008.48.

[26] G. Choijilsuren, R.-S. Jhou, S.-F. Chou, C.-J. Chang, H.-I. Yang, Y.-Y. Chen, W.-L. Chuang, M.-L. Yu, C. Shih, Heparin at physiological concentration can enhance PEG-free in vitro infection with human hepatitis B virus, Sci. Rep. 7 (2017) 14461. 10.1038/s41598-017-14573-9.

[27] M.A. Lovich, E.R. Edelman, Tissue concentration of heparin, not administered dose, correlates with the biological response of injured arteries in vivo, Proc. Natl. Acad. Sci. U. S. A. 96 (1999) 11111–11116.

[28] M. Mitsi, K. Forsten-Williams, M. Gopalakrishnan, M.A. Nugent, A Catalytic Role of Heparin within the Extracellular Matrix, J. Biol. Chem. 283 (2008) 34796–34807. 10.1074/jbc.M806692200.

[29] G. Carpentier, S. Berndt, S. Ferratge, W. Rasband, M. Cuendet, G. Uzan, P. Albanese, Angiogenesis Analyzer for ImageJ — A comparative morphometric analysis of “Endothelial Tube Formation Assay” and “Fibrin Bead Assay,” Sci. Rep. 10 (2020) 11568. 10.1038/s41598-020-67289-8.

[30] M. Ohlin, A new look at a poorly immunogenic neutralization epitope on cytomegalovirus glycoprotein B. Is there cause for antigen redesign?, Mol. Immunol. 60 (2014) 95–102. 10.1016/j.molimm.2014.03.015.

[31] X. Wang, J.A. Maynard, The Bordetella Adenylate Cyclase Repeat-in-Toxin (RTX) Domain Is Immunodominant and Elicits Neutralizing Antibodies, J. Biol. Chem. 290 (2015) 3576– 3591. 10.1074/jbc.M114.585281.

[32] W.-C. Liang, X. Wu, F.V. Peale, C.V. Lee, Y.G. Meng, J. Gutierrez, L. Fu, A.K. Malik, H.-P. Gerber, N. Ferrara, G. Fuh, Cross-species Vascular Endothelial Growth Factor (VEGF)-blocking Antibodies Completely Inhibit the Growth of Human Tumor Xenografts and Measure the Contribution of Stromal VEGF *, J. Biol. Chem. 281 (2006) 951–961. 10.1074/jbc.M508199200.

[33] G. Fuh, P. Wu, W.-C. Liang, M. Ultsch, C.V. Lee, B. Moffat, C. Wiesmann, Structure-function studies of two synthetic anti-vascular endothelial growth factor Fabs and comparison with the Avastin Fab, J. Biol. Chem. 281 (2006) 6625–6631. 10.1074/jbc.M507783200.

[34] R. Mamluk, Z. Gechtman, M.E. Kutcher, N. Gasiunas, J. Gallagher, M. Klagsbrun, Neuropilin-1 Binds Vascular Endothelial Growth Factor 165, Placenta Growth Factor-2, and Heparin via Its b1b2 Domain *, J. Biol. Chem. 277 (2002) 24818–24825. 10.1074/jbc.M200730200.

[35] K.L. DeCicco-Skinner, G.H. Henry, C. Cataisson, T. Tabib, J.C. Gwilliam, N.J. Watson, E.M. Bullwinkle, L. Falkenburg, R.C. O’Neill, A. Morin, J.S. Wiest, Endothelial Cell Tube Formation Assay for the In Vitro Study of Angiogenesis, J. Vis. Exp. JoVE (2014) 51312. 10.3791/51312.

[36] Y. Liu, H. Tian, G.C. Blobe, C.P. Theuer, H.I. Hurwitz, A.B. Nixon, Effects of the combination of TRC105 and bevacizumab on endothelial cell biology, Invest. New Drugs 32 (2014) 851–859. 10.1007/s10637-014-0129-y.

[37] L. Xiang, R. Varshney, N.A. Rashdan, J.H. Shaw, P.G. Lloyd, Placenta Growth Factor and Vascular Endothelial Growth Factor-A Have Differential, Cell-Type Specific atterns of Expression in Vascular Cells, Microcirc. N. Y. N 1994 21 (2014) 368–379. 10.1111/micc.12113.

[38] A. Luttun, K. Brusselmans, H. Fukao, M. Tjwa, S. Ueshima, J.-M. Herbert, O. Matsuo, D. Collen, P. Carmeliet, L. Moons, Loss of placental growth factor protects mice against vascular permeability in pathological conditions, Biochem. Biophys. Res. Commun. 295 (2002) 428–434. 10.1016/s0006-291x(02)00677-0.

[39] H. Bessho, B. Wong, D. Huang, E.Y. Siew, D. Huang, J. Tan, C.K. Ong, S.Y. Tan, K. Matsumoto, M. Iwamura, B.T. Teh, Inhibition of Placental Growth Factor in Renal Cell Carcinoma, ANTICANCER Res. (2015).

[40] T.-H. Lee, S. Seng, M. Sekine, C. Hinton, Y. Fu, H.K. Avraham, S. Avraham, Vascular Endothelial Growth Factor Mediates Intracrine Survival in Human Breast Carcinoma Cells through Internally Expressed VEGFR1/FLT1, PLOS Med. 4 (2007) e186. 10.1371/journal.pmed.0040186.

[41] N. Al-Zeheimi, Y. Gao, P.A. Greer, S.A. Adham, Neuropilin-1 Knockout and Rescue Confirms Its Role to Promote Metastasis in MDA-MB-231 Breast Cancer Cells, Int. J. Mol. Sci. 24 (2023) 7792. 10.3390/ijms24097792.

[42] A.M. Devery, R. Wadekar, S.M. Bokobza, A.M. Weber, Y. Jiang, A.J. Ryan, Vascular endothelial growth factor directly stimulates tumour cell proliferation in non-small cell lung cancer, Int. J. Oncol. 47 (2015) 849–856. 10.3892/ijo.2015.3082.

[43] A. Nasir, L.O. Reising, D.M. Nedderman, A.D. Fulford, M.T. Uhlik, L.E. Benjamin, A.E. Schade, T.R. Holzer, Heterogeneity of Vascular Endothelial Growth Factor Receptors 1, 2, 3 in Primary Human Colorectal Carcinoma, Anticancer Res. 36 (2016) 2683–2696.

[44] M. Luo, L. Hou, J. Li, S. Shao, S. Huang, D. Meng, L. Liu, L. Feng, P. Xia, T. Qin, X. Zhao, VEGF/NRP-1axis promotes progression of breast cancer via enhancement of epithelial-mesenchymal transition and activation of NF-κB and β-catenin, Cancer Lett. 373 (2016) 1– 11. 10.1016/j.canlet.2016.01.010.

[45] M.T. Dellinger, R.A. Brekken, Phosphorylation of Akt and ERK1/2 Is Required for VEGF-A/VEGFR2-Induced Proliferation and Migration of Lymphatic Endothelium, PLoS ONE 6 (2011) e28947. 10.1371/journal.pone.0028947.

[46] T. Zhong, L. Zhang, Z. Huang, X. Pang, C. Jin, W. Liu, J. Du, W. Yin, N. Chen, J. Min, M. Xia, B. Li, Design of a fragment crystallizable-engineered tetravalent bispecific antibody targeting programmed cell death-1 and vascular endothelial growth factor with cooperative biological effects, iScience 28 (2025) 111722. 10.1016/j.isci.2024.111722.

[47] D. Krilleke, A. DeErkenez, W. Schubert, I. Giri, G.S. Robinson, Y.-S. Ng, D.T. Shima, Molecular mapping and functional characterization of the VEGF164 heparin-binding domain, J. Biol. Chem. 282 (2007) 28045–28056. 10.1074/jbc.M700319200.

[48] N. Haspel, D. Zanuy, R. Nussinov, T. Teesalu, E. Ruoslahti, C. Aleman, Binding of a C-end rule peptide to neuropilin-1 receptor: A molecular modeling approach, Biochemistry 50 (2011) 1755–1762. 10.1021/bi101662j.

[49] H. Rouha, A. Badarau, Z.C. Visram, M.B. Battles, B. Prinz, Z. Magyarics, G. Nagy, I. Mirkina, L. Stulik, M. Zerbs, M. Jägerhofer, B. Maierhofer, A. Teubenbacher, I. Dolezilkova, K. Gross, S. Banerjee, G. Zauner, S. Malafa, J. Zmajkovic, S. Maier, R. Mabry, E. Krauland, K.D. Wittrup, T.U. Gerngross, E. Nagy, Five birds, one stone: Neutralization of α-hemolysin and 4 bi-component leukocidins of Staphylococcus aureus with a single human monoclonal antibody, mAbs 7 (2014) 243–254. 10.4161/19420862.2014.985132.

