## Supplemental files for "Antibodies blocking PlGF or VEGF interactions with the NRP1 receptor mediate anti-proliferative effects"

[a]McKetta Department of Chemical Engineering
University of Texas at Austin
200 E Dean Keeton St
Austin, TX 78712

[b]Interdisciplinary Life Sciences Graduate Program
The University of Texas at Austin
North Hackerman Building
100 East 24th St, NHB 4500
Austin, TX 78712

[c]Department of Chemical and Biological Engineering
University of Colorado Boulder
3415 Colorado Ave
Boulder, CO 80309, USA

[d]University of Hawaii Comprehensive Cancer Center
University of Hawaii
701 Ilao St
Honolulu, HI 96813, USA

[e]AO Research Institute Davos
Clavadelerstrasse 8,
7270 Davos Platz, Switzerland

**Supplementary methods: qPCR**

To validate HUVEC gene expression in tube formation assay on Matrigel, we completed quantitative gene expression for VEGF, PlGF, NRP1, VEGFR1, and VEGFR2. HUVECs were seeded 5000 cell/well on Matrigel and let incubate for 16-18 h, just like the tube formation assay. 16 wells per replicate were combined for adequate RNA collection. RNA was then isolated by RNeasy mini kit (Qiagen) and converted to cDNA by SuperScript VILO (ThermoFisher). The following primer/probe TaqMan™ Assays (ThermoFisher) were used for gene identification on a QuantStudio™ Pro6 (ThermoFisher): VEGFA (HS00900055_m1), PlGF (HS00182176), NRP1 (HS00826128_m1), VEGFR1 (HS00176573_m1), and VEGFR2 (HS00176676_m1).Relative gene expression was calculated by the following formula, assuming 100% primer efficiency, where the $E_{ref}$ was the house keeping gene RPLP0 (MicroSynth):

$$RE = \frac{\left(E_{Ref}\right)^{Ct_{Ref}}}{\left(E_{GOI}\right)^{Ct_{GOI}}}$$

### A, Human PLGF1 vs PlGF2

```
huPlGF1        LPAVPPQQWALSAGNGSSEVEVVPFQEVWGRSYCRALERLVDVVSEYPSEVEHMFSPSCV   60
huPlGF2        LPAVPPQQWALSAGNGSSEVEVVPFQEVWGRSYCRALERLVDVVSEYPSEVEHMFSPSCV   60
               ************************************************************

huPlGF1        SLLRCTGCCGDENLHCVPVETANVTMQLLKIRSGDRPSYVELTFSQHVRCECRPLREKMK   120
huPlGF2        SLLRCTGCCGDENLHCVPVETANVTMQLLKIRSGDRPSYVELTFSQHVRCECRPLREKMK   120
               ************************************************************

huPlGF1        PER---------------------CGDAVPRR   131
huPlGF2        PERRRPKGRGKRRREKQRPTDCHLCGDAVPRR   152
```

### B, Human PlGF2 vs VEGFA$_{165}$

```
VEGF-165       --------APMAEGGGQNHHEVVKFMDVYQRSYCHPIETLVDIFQEYPDEIEYIFKPSCV   52
huPlGF2        LPAVPPQQWALSAGNGSSEVEVVPFQEVWGRSYCRALERLVDVVSEYPSEVEHMFSPSCV   60
                         :: *.*... *** * :*: ****: :* ***:..***.*:*::*.****

VEGF-165       PLMRCGGCCNDEGLECVPTEESNITMQIMRIKPHQGQHIGEMSFLQHNKCECRPKKDRAR   112
huPlGF2        SLLRCTGCCGDENLHCVPVETANVTMQLLKIRSGDRPSYVELTFSQHVRCECRPLREKMK   120
                *:** ***.**.*.***.* :*:***:::*:  :     *::* ** :***** ::: :

VEGF-165       QEN--PCGPCSERRKHLFVQDPQTCKCSCKNTDSRCKARQLELNERTCRCDKPRR   165
huPlGF2        PERRRPKGRGKRRRE---KQRPTDCHLCGD--------------------AVPRR   152
                *.  * *  ..**:    * *  *: . .                      ***
```

### C, Human PlGF2 vs mouse PlGF2

```
huPlGF2        LPAVPPQQWALSAGNGSSEVEVVPFQEVWGRSYCRALERLVDVVSEYPSEVEHMFSPSCV   60
mPlGF2         ----VHSQGALSAGNNSTEVEVVPFNEVWGRSYCRPMEKLVYILDEYPDEVSHIFSPSCV   56
                     .* ******.*:*******:********* :*:** ::.***.**.*:******

huPlGF2        SLLRCTGCCGDENLHCVPVETANVTMQLLKIRSGDRP-SYVELTFSQHVRCECRPLREKM   119
mPlGF2         LLSRCSGCCGDEGLHCVPIKTANITMQILKIPPNRDPHFYVEMTFSQDVLCECRPILETT   116
                * **:******.*****::***:***:***   . *  ***:****.* *****: *.

huPlGF2        KPERRRPKGRGKRRREKQRPTDCHLCGDAVPRR   152
mPlGF2         KAERRKTKGKRKRSRNSQTEEPHP---------   140
                * ***: **: ** *:.*
```

### D, Human VEGFA$_{121}$ vs VEGFA$_{165}$

```
VEGF-121       APMAEGGGQNHHEVVKFMDVYQRSYCHPIETLVDIFQEYPDEIEYIFKPSCVPLMRCGGC   60
VEGF165        APMAEGGGQNHHEVVKFMDVYQRSYCHPIETLVDIFQEYPDEIEYIFKPSCVPLMRCGGC   60
                ************************************************************

VEGF-121       CNDEGLECVPTEESNITMQIMRIKPHQGQHIGEMSFLQHNKCECRPKKDRARQEK-----   120
VEGF165        CNDEGLECVPTEESNITMQIMRIKPHQGQHIGEMSFLQHNKCECRPKKDRARQENPCGPC   120
                ******************************************************:

VEGF-121       ---------------------------------------CDKPRR      121
VEGF165        SERRKHLFVQDPQTCKCSCKNTDSRCKARQLELNERTCRCDKPRR      165
```

**Supplemental Fig. S1. Sequence alignments of growth factor isoforms. A,** Human PlGF1 (Uniprot P49763-2) versus human PlGF2 (Uniprot P49763-3). The PlGF2 exon 6 insert that encodes the heparin binding domain is highlighted in yellow, the heparin binding motifs (BBXB) are underlined, the c-terminal NRP1-binding peptide highlighted in blue and cysteines in bold. **B,** Comparison of the 42% identical human PlGF2 versus

VEGFA165 (Uniprot P15692-4) sequences, with conserved cysteines in bold and putative or confirmed heparin binding sites underlined (R123, R124, R159 for VEGF). **C,** Comparison of the 60% identical human PlGF2 vs mouse PlGF2 (Uniprot # Q544A5), **D,** VEGFA165 versus VEGFA121 (Uniprot P15692-9) showing the exon 7 and 8a sequences responsible for heparin- and NRP1-binding, respectively. Alignments performed with Clustal Omega.

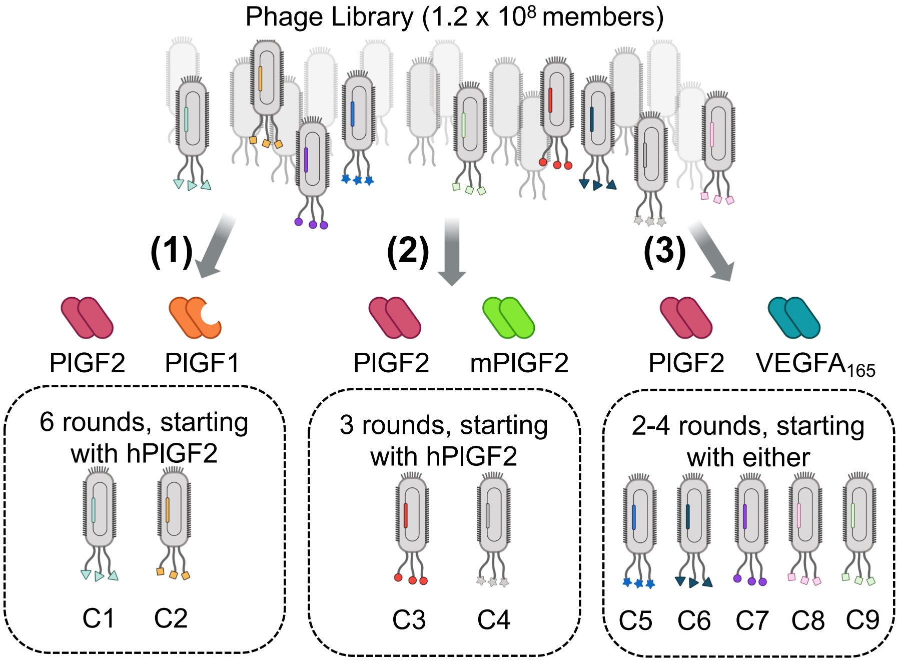


**Supplemental Figure S2. Phage panning strategies.** A phage library was generated from mice immunized with hPlGF2 and panned using three strategies: **(1)** hPlGF2 with screening to identify clones not binding PLGF1, **(2)** alternate panning against mPlGF2 and hPlGF2, and **(3)** alternate panning against hVEGFA165 and hPlGF2 to yield the indicated clones.

**A.**

| Clone | $V_H$ germline chain usage | | | $V_H$ somatic mutations | $V_L$ germline chain usage | | $V_L$ somatic mutations |
|---|---|---|---|---|---|---|---|
| ***C1*** | IGHV14-4 | IGHD2-4 | IGHJ4 | 2 | IGKV6-15 | IGKJ2 | 5 |
| ***C2*** | IGHV1-9 | IGHD1-1 | IGHJ3 | 16 | IGKV6-15 | IGKJ5 | 9 |
| ***C3*** | IGHV1-47 | IGHD6-1 | IGHJ4 | 10 | IGKV6-17 | IGKJ1 | 2 |
| ***C4*** | IGHV1-9 | IGHD2-4 | IGHJ2 | 15 | IGKV4-57 | IGKJ5 | 6 |
| ***C5*** | IGHV1-47 | IGHD2-2 | IGHJ4 | 6 | IGKV12-44 | IGKJ5 | 4 |
| ***C6*** | IGHV14-4 | IGHD4-1 | IGHJ4 | 6 | IGKV3-4 | IGKJ1 | 4 |
| ***C7*** | IGHV14-4 | IGHD1-1 | IGHJ2 | 3 | IGKV6-15 | IGKJ2 | 2 |
| ***C8*** | IGHV1-9 | IGHD2-14 | IGHJ4 | 2 | IGKV14-126 | IGKJ5 | 4 |
| ***C9*** | IGHV1-18 | IGHD6-10 | IGHJ4 | 5 | IGKV3-7 | IGKJ2 | 3 |

**B.**

---

>G6-31 VH
EVQLVESGGGLVQPGGSLRLSCAASG**FTISDYWIH**WVRQAPGKGLEWVAG**ITPAGGYTYYADSVKG**RFTISADTSKN TAYLQMNSLRAEDTAVYYCAR**FVFFLPYAMDY**WGQGTLV

>G6-31 VL

DIQMTQSPSSLSASVGDRVTITC**RASQDVSTAVA**WYQQKPGKAPKLLIY**SASFLYS**GVPSRFSGSGSGTDFTLTISSLQPE DFATYYC**QQGYGNP**FTFGQGTKVEIKR

---

>C9.V2 VH
EVQLVESGGGLVQPGGSLRLSCAASG**FTFTNTWIS**WVRQAPGKGLEWVGS**ITPANGYTNYADSVKG**RFTISADTSKN TAYLQMNSLRAEDTAVYYCAR**AVYPWFFAY**WGQGTLVTVS

>C9.V2 VL

DIQMTQSPSSLSASVGDRVTITC**RASQYVSHAVA**WYQQKPGKAPKLLIY**SASFLYS**GVPSRFSGSGSGTDFTLTISSLQP EDFATYYC**QSAYTPPTT**FGQGTKVEIKR

---

>C1 VH
EVQLQQSGAELVRSGASVKLSCTASG**FNIKDYYMH**WVKQRPEQGLEWIGW**IDPEDGDTEYAPKFQG**KATMTADTSS NTAYLQLSSLTSEDTAVYYCNA**PDDDYDSSYAMDY**WGQGTSVTVS

>C1 VL

DIVMTQSQKFMSTSVGDRVSVTC**KASQNVGTNVA**WYQQKPGQSPKALIY**SASYRYS**GVPDRFTGSGSGTDFTLTISNV QSEDLAEYIC**QQYDTFPYT**FGGGTKLEIKR

---

>C2 VH

QVQLQQSGAELMKPGASVKISCKAPG**YSFTSYWIE**WVKQRPGHGLEWIGE**ILPGSISPNYNEQFRG**KATITADISSNT AYMQLSSLTSEDSAVYYCAR**ERAVYGSKFAY**WGQGTLVTVS

>C2 VL

DIVMTQSHKFMSASVGDRVSITC**KASQDVGTAVA**WYQQKPGQSPKALIY**SASYRYS**GVPDRFTGSGSGTDFTLTISNVQSEDLAEYFC**QQYNSYPLT**FGAGTKLELKR

---

>C3 VH

QVQLQQPGAELVRPGASVKLSCKASG**YTFTSYWIN**WVKQRPGQGLEWIGN**IYPSDSYTNYNQKFRD**KATLTVDKSSSTAYMQLSRPTSEDSAVYYCSR**WDAPYAMDY**WGQGTSVTVS

>C3 VL

DIVMTQSHKFMSTSVGDRVSITC**KASQDVSTAVA**WYQQKPGQSPKLLIY**SASYRYT**GVPDRFTGSGSGTDFTFTISSVQAEDLAVYYC**QQHYSTPWT**FGGGTKLEIKR

---

>C4 VH

EVQLQQSGAELARPGASVKMSCKATG**YTFSIYWIE**WVKQRPGHGLEWIGE**ILPGSGSTNYNEKFKG**KATFTADTSSNTAYMQLSSLTSEDSAVYYCAR**WPIMIAGHYLDY**WGQGTTLTVSS

>C4 VL

DIVMTQSPAIMSASPGEKVTITC**SASSSVSYMH**WFQQKPGTSPKLWIY**STSNLAS**GVPARFSGSGSGTSYSLTISRMEAEDAATYYC**QQRSSYPLT**FGAGTKLEIKR

---

>C5 VH

QVQLQQSGAELVRPGASVKLSCKASG**YTFTSYWIN**WVKQRPGQGLEWIGN**IYPSDGYTNYNQKFKD**KATLTVDKSTSTAYMQLSSPTSEDSAVYYCTR**WYYAYGGGFAMDY**WGQGTSVTVS

>C5 VL

DIVMTQSPASLSASVGETVTITC**RASENIYSYLA**WYQQKQGKSPQLLVY**NAKTLAE**GVPSRFSGSGSGTQFSLKINSLQPEDFGSYYC**QHHYGTPLT**FGAGTKLELKR

---

>C6 VH

DVKLRESGAELVRSGASVKLSCTASG**FNIKDYYMH**WVKQRPEQGLEWIGW**IDPENGDTEYAPKFQG**KATMTADTSSNTAYLQLSSLTSEDTAVYYCNA**WGAMDY**WGQGTSVTVS

>C6 VL

DIVMTQSPASLAVSLGQRATISC**KASQSVDYDGDSYMN**WYQQKPGQPPKLLIY**AASNLES**GIPARFSGSGSGTDFTLNIHPVEEEDAATYYC**QQSYEDPWT**FGGGTKLEIKR

---

>C7 VH

QVQLQQSGAELVRSGASVKLSCTASG**FNIKDYYMH**WVKQRPEQGLEWIGW**IDPENGDTEYAPKFQG**KATMTADTSSNTAYLQLSSLTSEDTAVYYCNA**WYYGSSYNYFDY**WGQGTTLTVS

>C7 VL

DIVMTQAQKFMSTSVGDRVSVTC**KASQNVGTNVA**WYQQKPGQSPKALIY**SASYRYS**GVPDRFTGSGSGTDFTLTNSNVQSEDLAEYFC**QQYNSYPPT**FGGGTKLEIKR

---

>C8 VH

QVQLQQSGAELMKPGASVKISCKATG**YTFSSYWIE**WVKQRPGHGLEWIGE**ILPGSGSTNYNEKFKG**KATFTADTSSNTAYMQLSSLTSEDSAVYYCAR**SYRYDDYAMDY**WGQGTSVTVS

>C8 VL

DIQMTQSPSSMYASLGERVTITC**KASQDIKSYLS**WYQQKPWKSPKTLIY**YATSLAD**GVPSRFSGSGSGQDYSLTISSLES
DDTATYYC**LQHGESPPT**FGAGTKLEIKR

---

>C9 VH

QVQLQQSGAELVRPGASVTLSCKASG**YTFTDYEMH**WAKQTPVHGLEWIGA**IDPETGGTAYNQNFKG**KATLTADKSSST
AYMELRSLTSEDSAVYYCAR**SYDYSAMDY**WGQGTSVTVS

>C9 VL

DIVMTQSPASLAVSLGQRATISC**RASQSVSTSSYSYMH**WYQQKPGQPPKLLIK**YASNLES**GAPARFSGSGSGTDFTLNI
HPVEEEDTATYYC**QHSWEIPYT**FGGGAKLEIKR

---

**Supplemental Figure S3. Antibody heavy and light chain sequences**. **A,** germline usage for each antibody. **B,** Sequences for the antibody variable region used in this work, with Kabat-defined CDRs underlined.

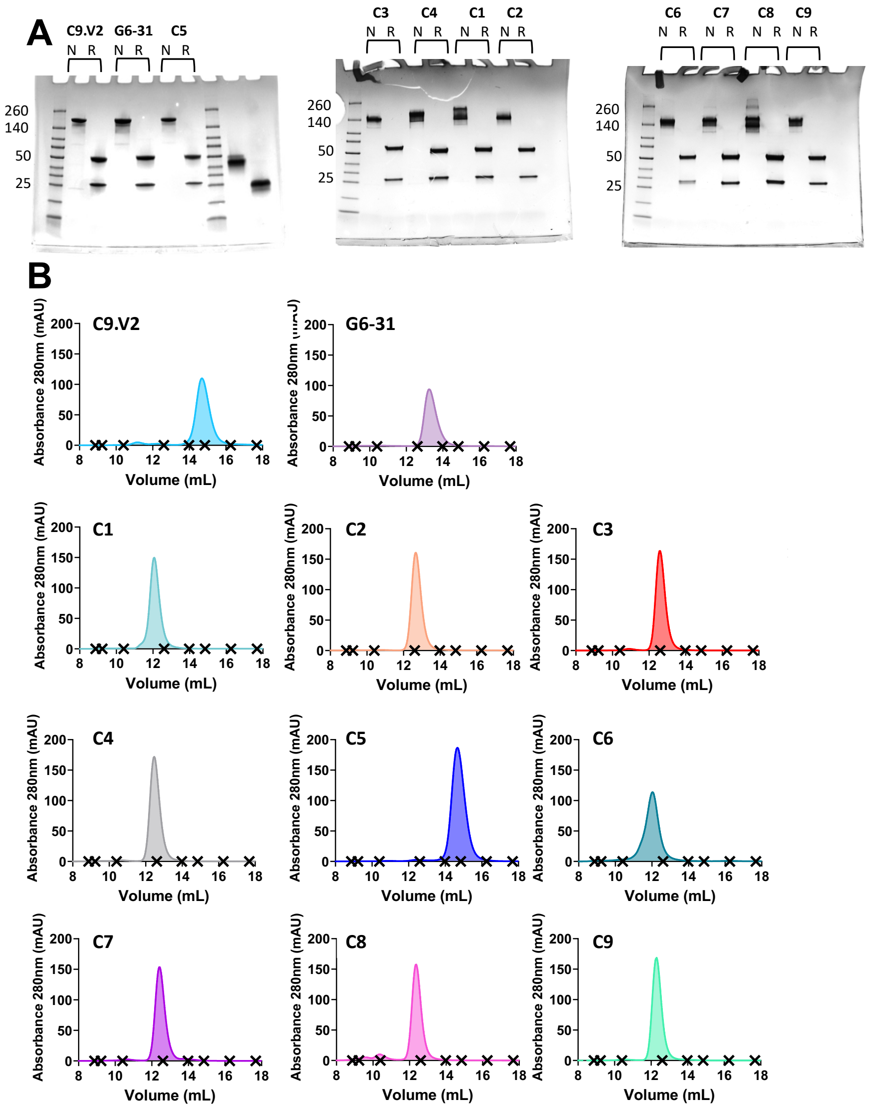


**Supplemental Figure S4. Antibody purity and monodispersity. A,** SDS-PAGE of purified antibodies with human IgG1/kappa domains. **B,** Size exclusion chromatography of on Superdex S200. 'X' icons indicate the observed elution volume for the following standard markers: (8.78 mL = 2000 kDa, 9.33 mL = 669 kDa, 10.55 mL = 440 kDa, 12.62 mL = 150 kDa, 14.02 mL =75 kDa, 14.89 mL = 44 kDa, 16.88 mL = 29 kDa, 17.7 mL =13.7 kDa).

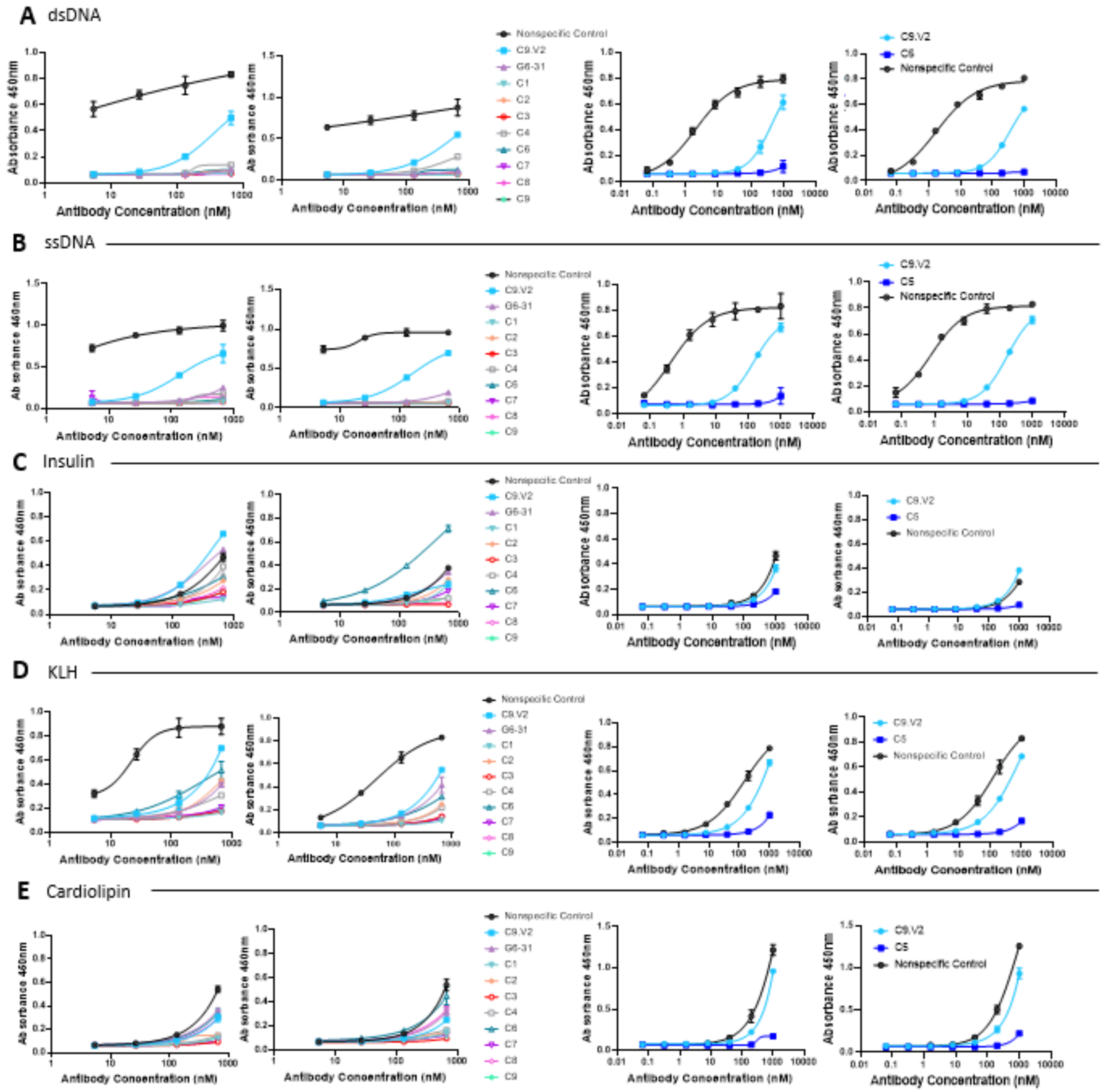


**Supplemental Figure S5.** Supplemental poly-specificity ELISAs (A) against double stranded DNA, (B) single stranded DNA, (C) insulin, (D) keyhole limpet hemocyanin from *Megathura crentulata* (KLH), and (E) cardiolipin. Duplicate ELISAs shown for each comparison; data shown are the mean and range of technical duplicates.

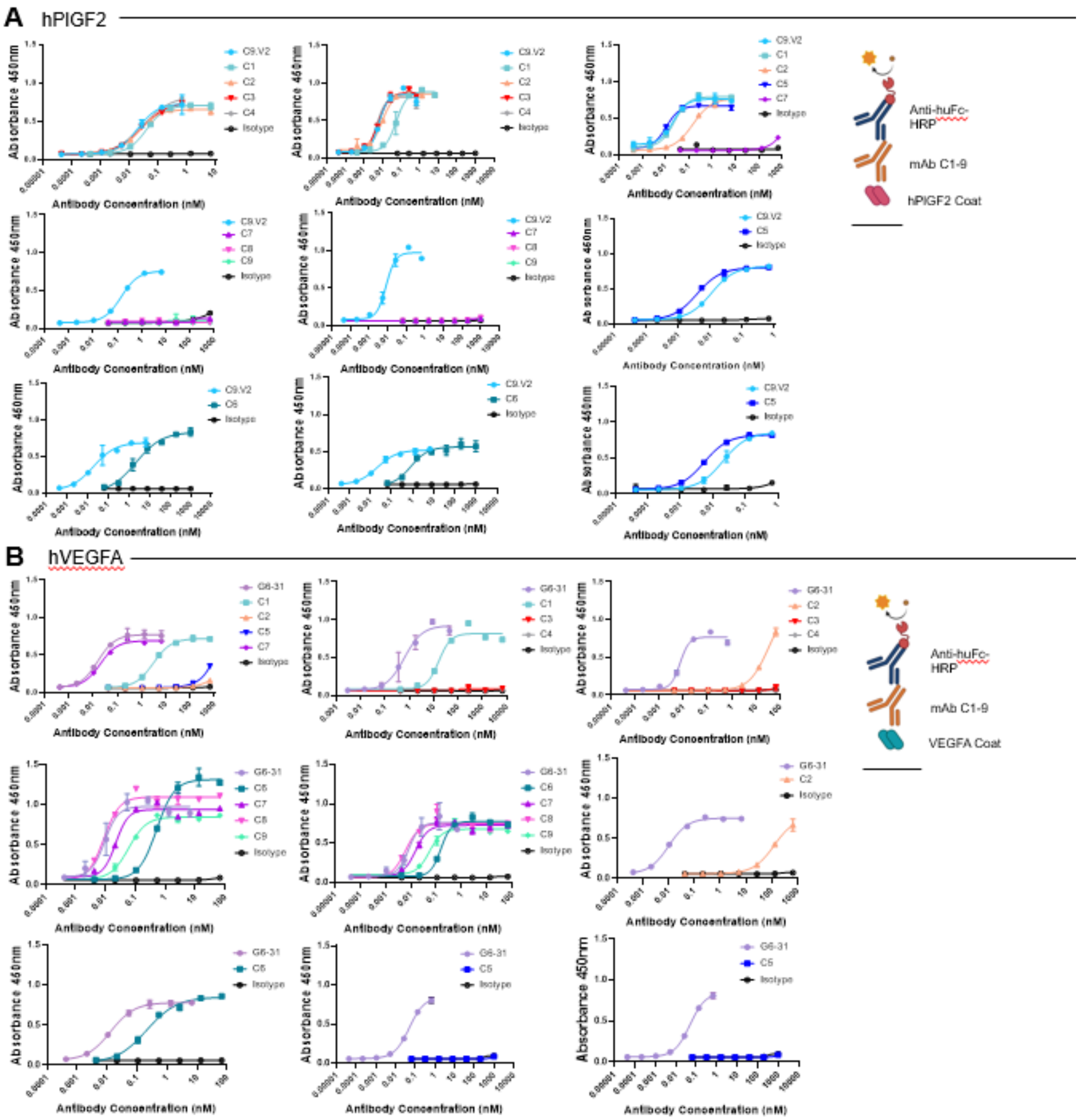


**Supplemental Figure S6.** ELISA data demonstrating PlGF2 and $VEGFA_{165}$ binding profiles. Replicate data shown for all antibodies binding to plates coated with (A) PlGF2, (B) $VEGFA_{165}$. Duplicate ELISAs shown for each comparison; data shown are the mean and range of technical duplicates.

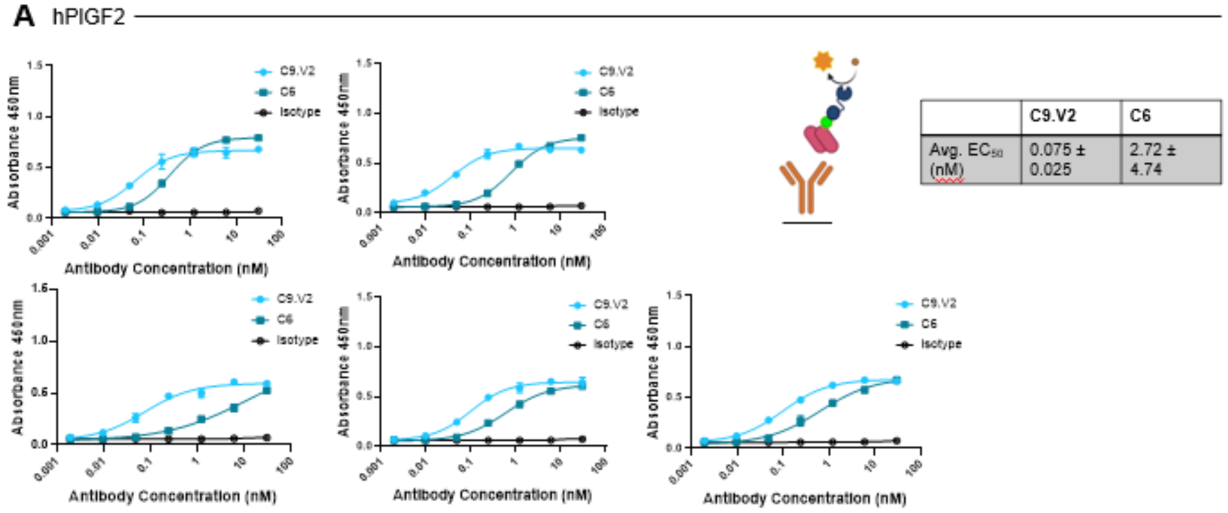


| | C9.V2 | C6 |
|---|---|---|
| Avg. $EC_{50}$ (nM) | 0.075 ± 0.025 | 2.72 ± 4.74 |

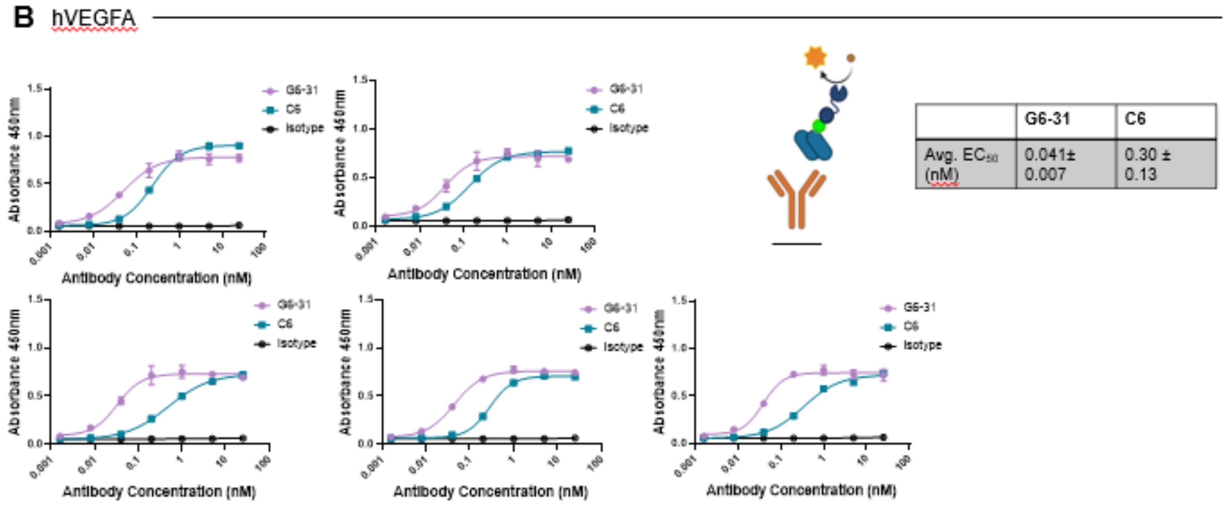


| | G6-31 | C6 |
|---|---|---|
| Avg. $EC_{50}$ (nM) | 0.041± 0.007 | 0.30 ± 0.13 |

**Supplemental Figure S7.** ELISA data demonstrates PIGF2 and $VEGFA_{165}$ binding profiles of cross-reactive antibody C6. These ELISAs were completed in a flipped orientation yo **Supp. Fig. 5**, where antibody was used to coat plate and growth factor was biotinylated and detected with streptavidin-HRP. Replicate data shown for all antibodies binding to plates coated with (A) PIGF2, (B) $VEGFA_{165}$. Duplicate ELISAs shown for each comparison; data shown are the mean and range of technical duplicates.

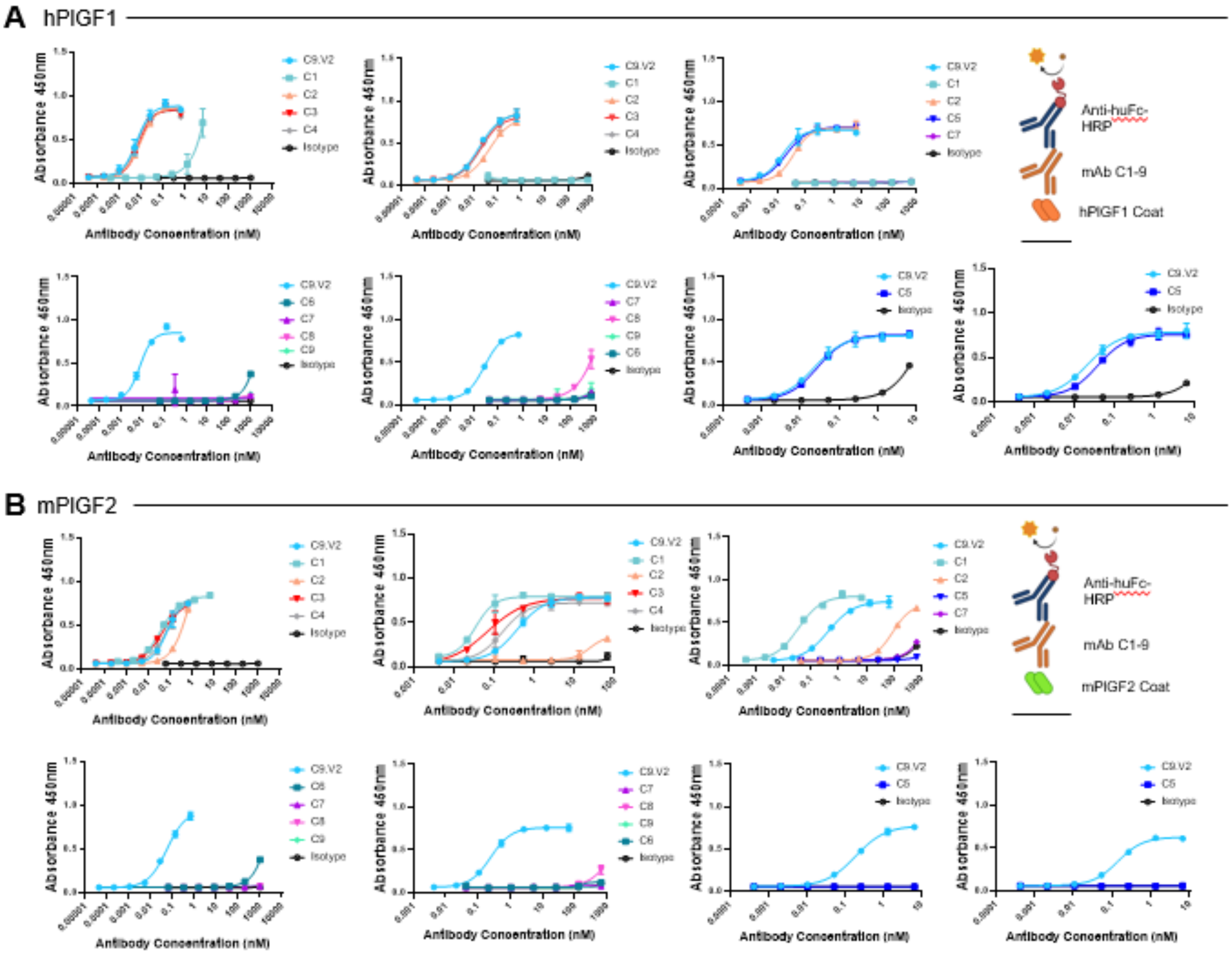


**Supplemental Figure S8.** ELISA data demonstrating PlGF2 and VEGFA$_{165}$ binding profiles. Replicate data shown for all antibodies binding to plates coated with (A) PlGF1, (B) mPlGF2. Duplicate ELISAs shown for each comparison; data shown are the mean and range of technical duplicates.

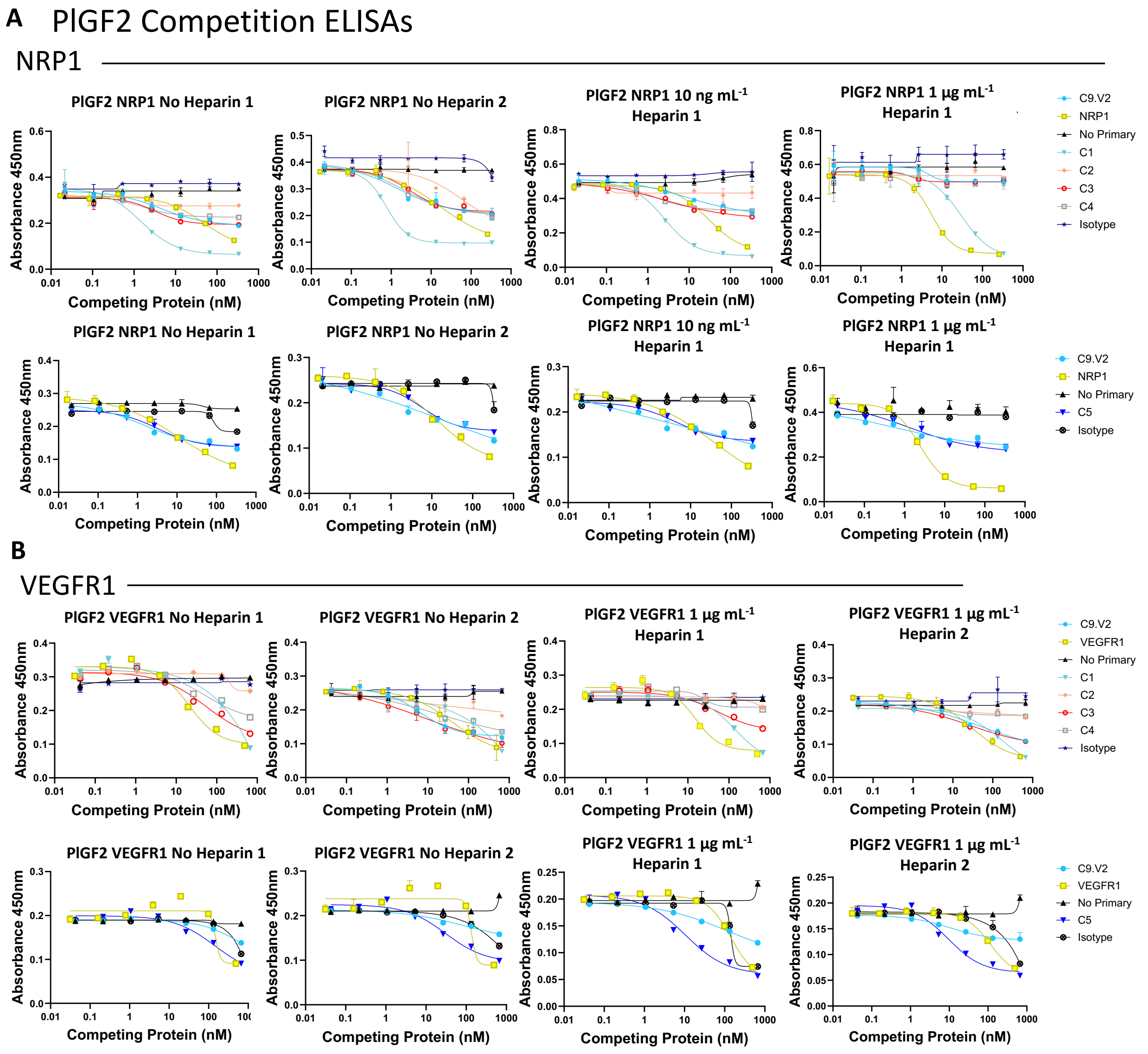


**Supplemental Information S9:** PlGF2 competition ELISAs of (A) antibodies versus NRP1 in duplicate and (B) antibodies versus VEGFR1. Data shown are the mean and range of technical duplicates.

### A VEGFA Competition ELISAs

#### NRP1

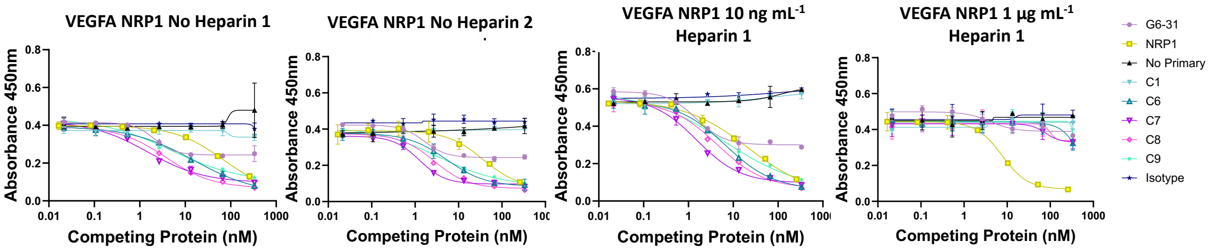


## B

#### VEGFR1

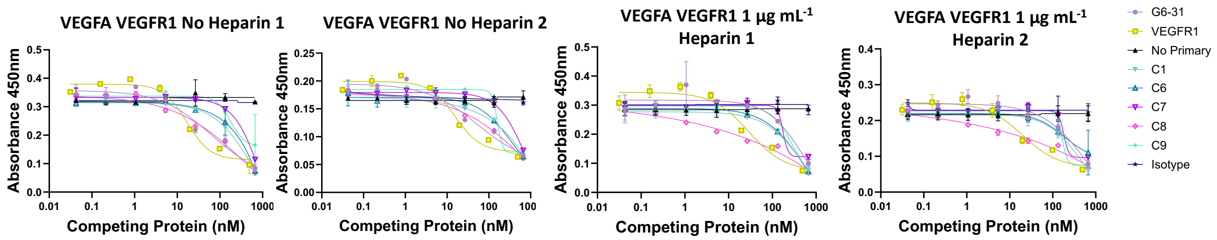


### C Pertinent IC50 values

IC50s of NRP1 Competition without heparin

| PIGF2 | NRP1 | C9.V2 | C1 | C5 |
|---|---|---|---|---|
| IC50 (nM) | 22.2 ± 17.6 | 6.5 ± 8.2 | 1.1 ± 0.5 | 4.5 1.7 |

| VEGFA | NRP1 | G6-31 | C6 | C7 | C8 | C9 |
|---|---|---|---|---|---|---|
| IC50 (nM) | 52.8 ± 22.1 | 1.9 ± 0.3 | 10.3 ± 7.0 | 1.4 ± 0.2 | 3.3 ± 0.7 | 4.1 ± 1.0 |

IC50s of VEGFR1 Competition without heparin

| PIGF2 | VEGFR1 | C9.V2 | C1 | C5 |
|---|---|---|---|---|
| IC50 (nM) | 141.7 ± 10.0 | 307.7 ± 304.1 | > 1000 | 89.7 ± 82.9 |

| VEGFA | VEGFR1 | G6-31 | C6 | C7 | C8 | C9 |
|---|---|---|---|---|---|---|
| IC50 (nM) | 16.3 ± 0.7 | 58.2 ± 6.9 | > 1000 | 975.7 ± 284.7 | 643.2 ± 440.8 | 168.2 ± 12.8 |

IC50s of VEGFR1 Competition with 1 µg/mL heparin

| PIGF2 | VEGFR1 | C9.V2 | C1 | C5 |
|---|---|---|---|---|
| IC50 (nM) | 75.1 ± 61.3 | 59.8 ± 55.9 | 155 ± 19.8 | 10.0 ± 0.7 |

**Supplemental Information S10:** $VEGFA_{165}$/ receptor competition ELISAs of (A) antibodies versus NRP1 in duplicate and (B) antibodies versus VEGFR1. Data shown are the mean and range of technical duplicates. (C) Pertinent $IC_{50}$ values from NRP1 and VEGFR1 competition for PlGF2 and $VEGFA_{165}$.

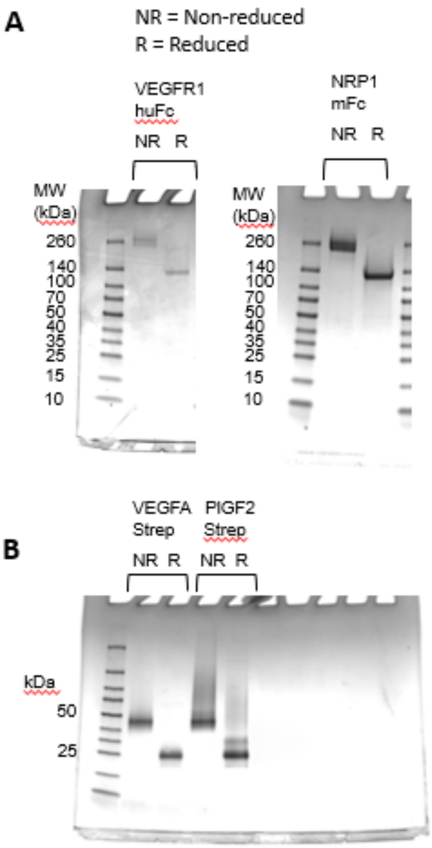


**Supplemental Information S11:** Production of recombinant receptors and growth factors. (A) SDS-PAGE of VEGFR1-huFc (203.6 kDa non-reduced, 101.8 kDa reduced) and NRP1-mFc (193 kDa non-reduced, 96.5 kDa reduced). (B) SDS-PAGE of VEGFA-2xStrep tagged (43.6 kDa non-reduced, 21.8 reduced) and PlGF2-2xStrep tagged (37.2 kDa non-reduced, 18.6 kDa reduced), purified by Streptactin resin.

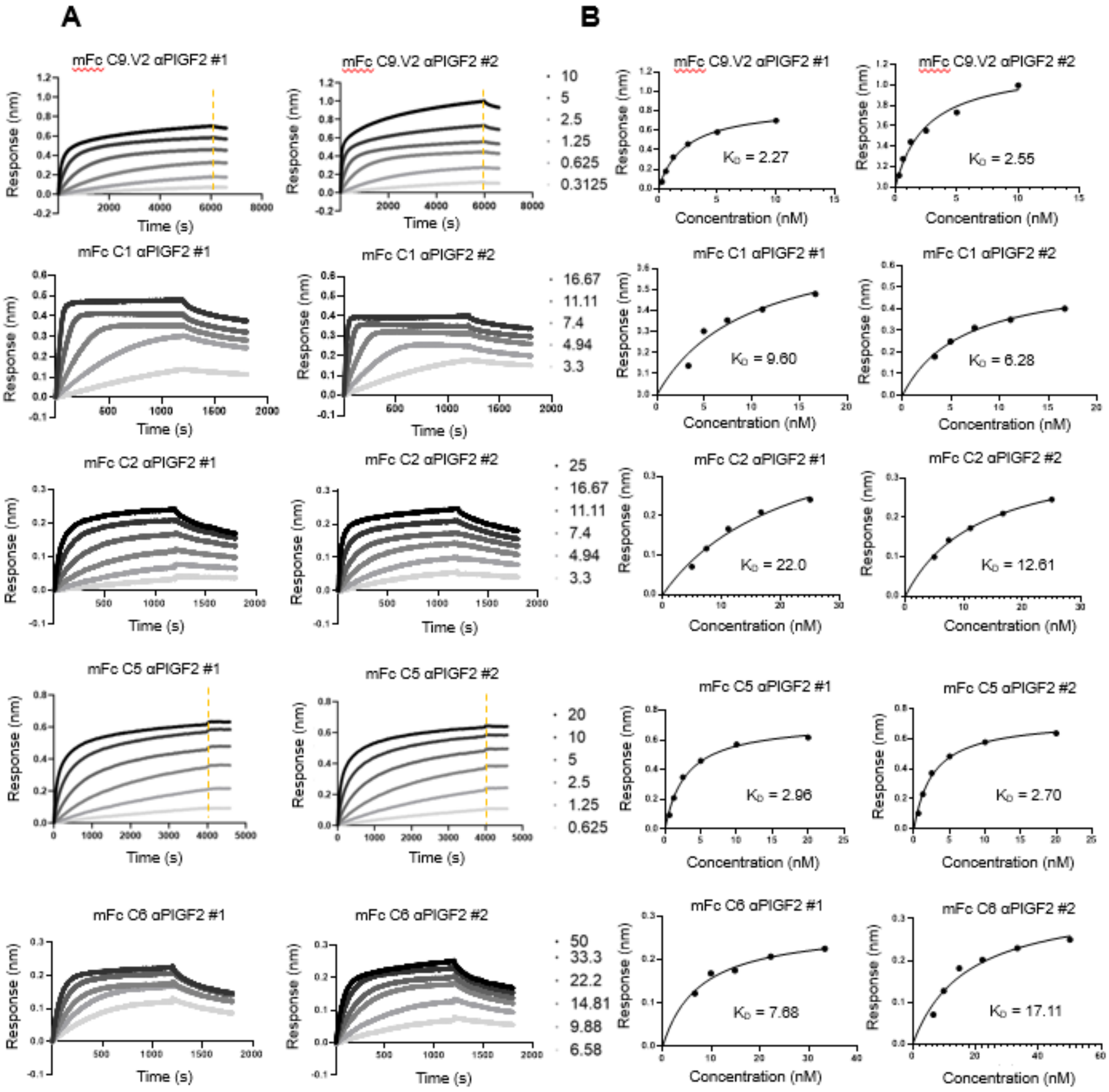


**Supplemental Information S12:** (A) BLI sensorgrams showing antibody binding to immobilized PlGF2, with technical duplicates. (B) The last ten association datapoints per dilution averaged and then fit to a one-site Langmuir isotherm to estimate effective $K_D$ values.

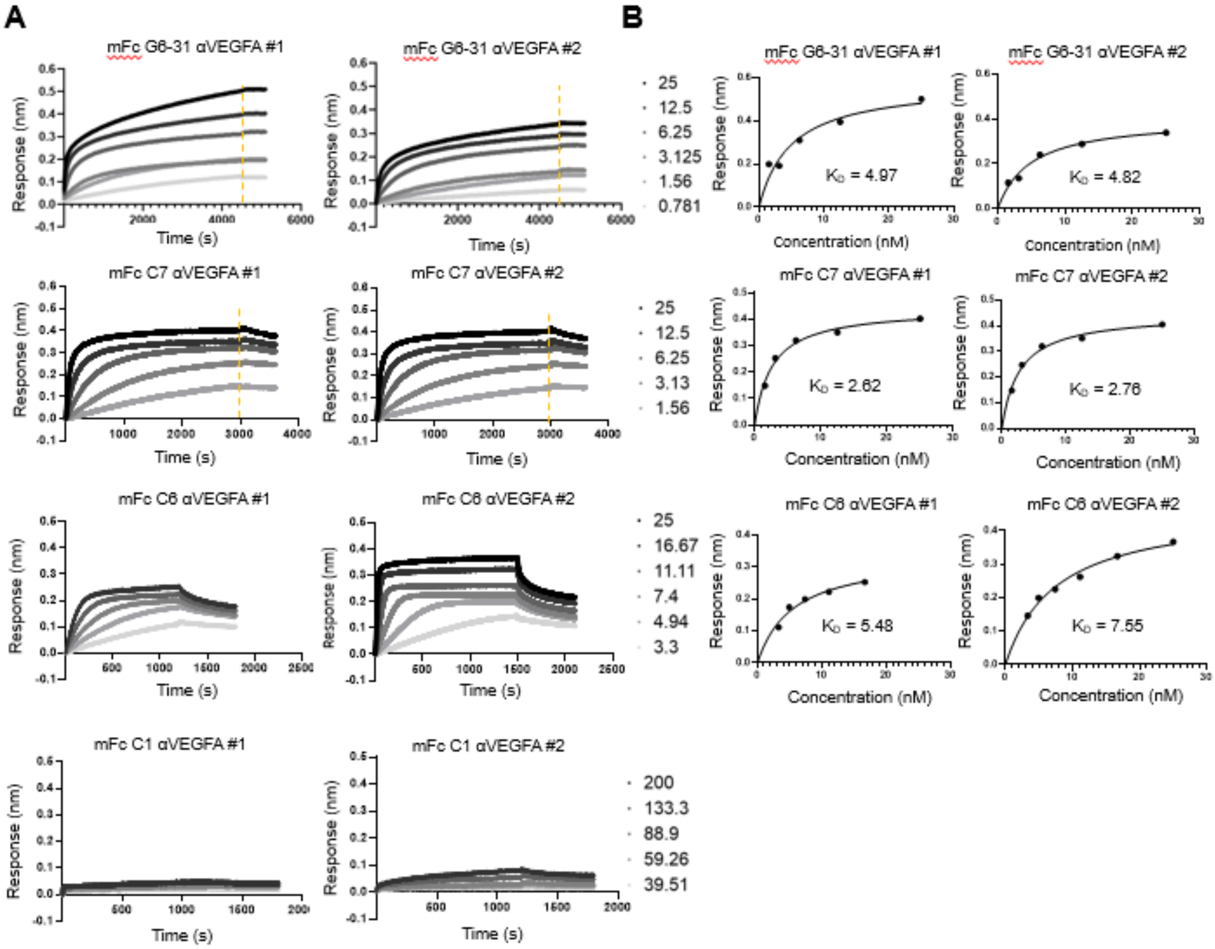


**Supplemental Information S13:** (A) BLI sensorgrams showing antibody binding to immobilized $VEGFA_{165}$, with technical duplicates. (B) The last ten association datapoints per dilution averaged and then fit to a one-site Langmuir isotherm.

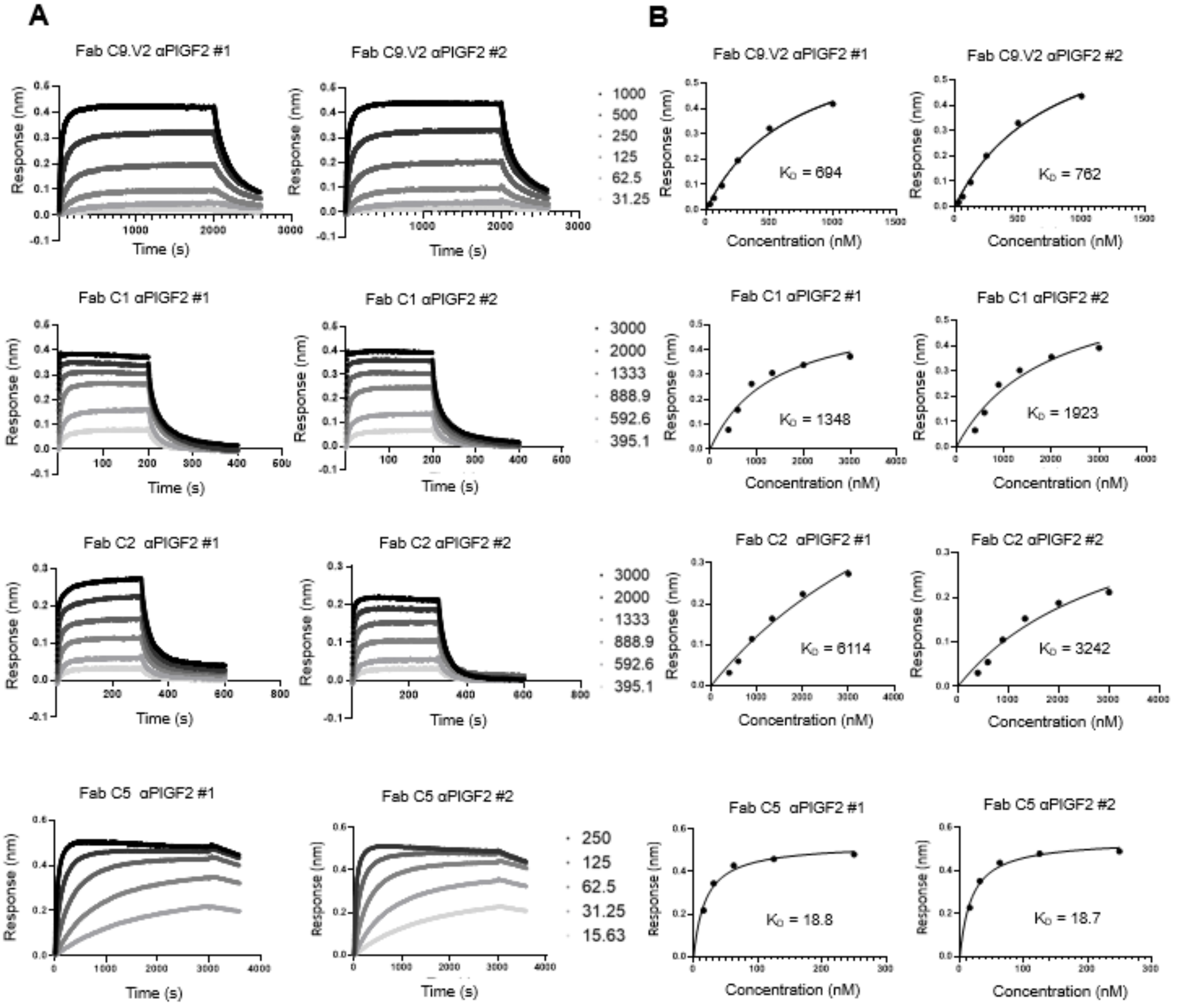


**Supplemental Information S14:** (A) Biolayer interferometry association and dissociation of Fabs to PlGF2, in duplicate. (B) The last ten association datapoints per dilution averaged and then fit to a one-site Langmuir isotherm.

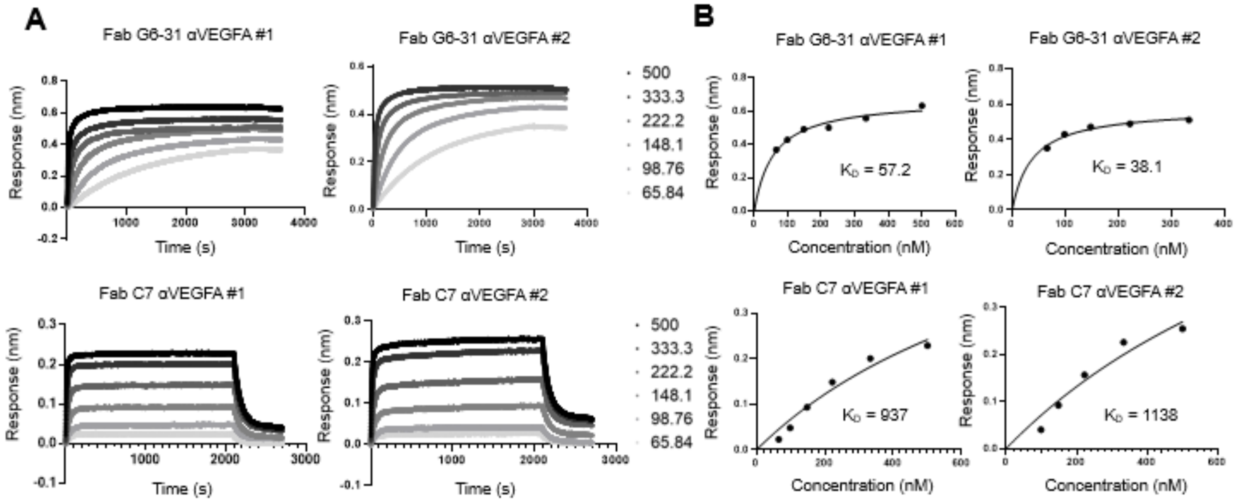


**Supplemental Information 14:** (A) Biolayer interferometry association and dissociation of Fabs to $VEGFA_{165}$, in duplicate. (B) The last ten association datapoints per dilution averaged and then fit to a one-site Langmuir isotherm.

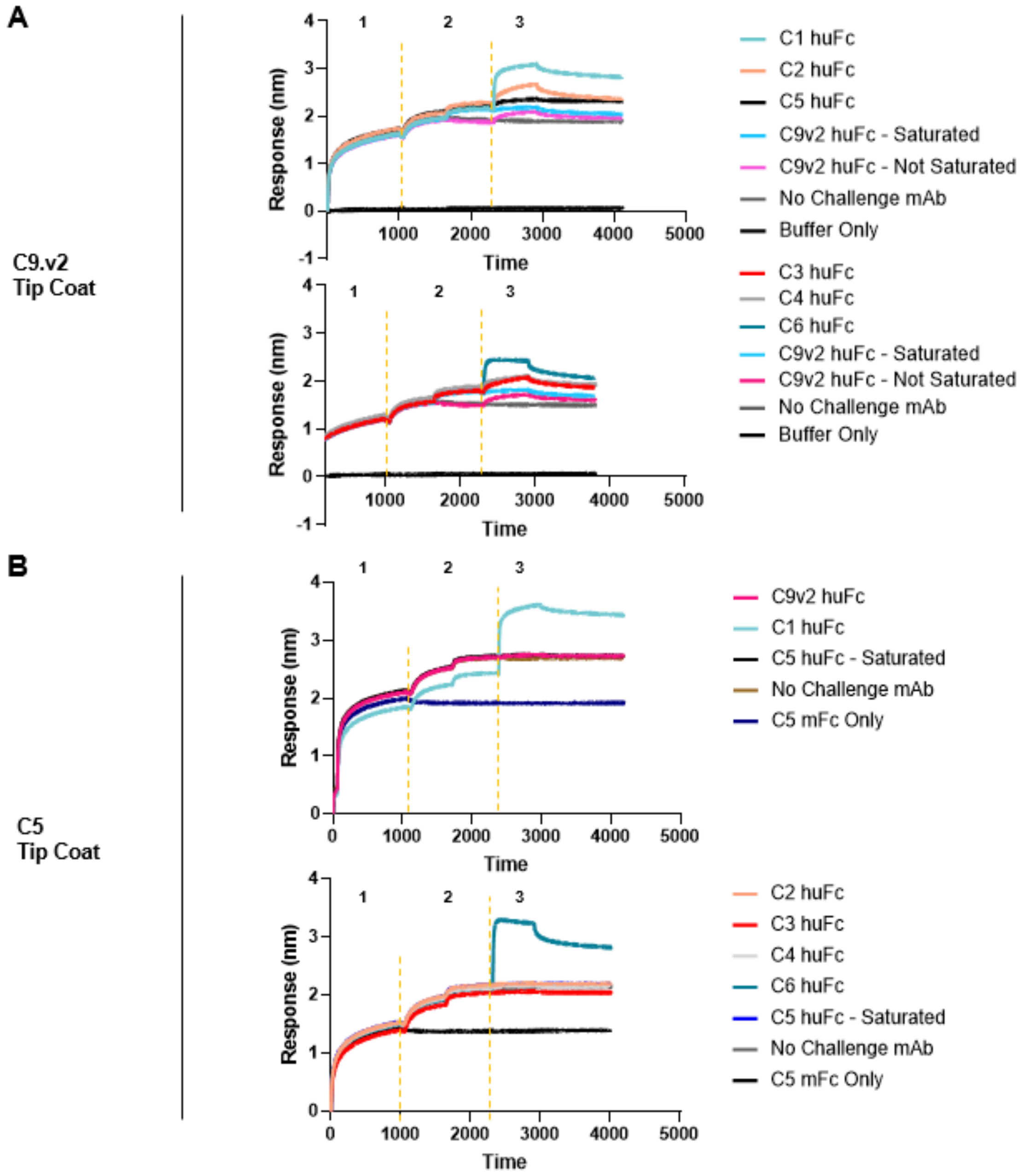


**Supplemental Information 15:** BLI responses used to bin anti-PlGF antibodies into epitope groups using (A) C9.V2 or (B) C5 as the capture antibody. Values shown in main Figure 3 were calculated by subtracting the signal at 500s from the signal at 0s of association (step 3).

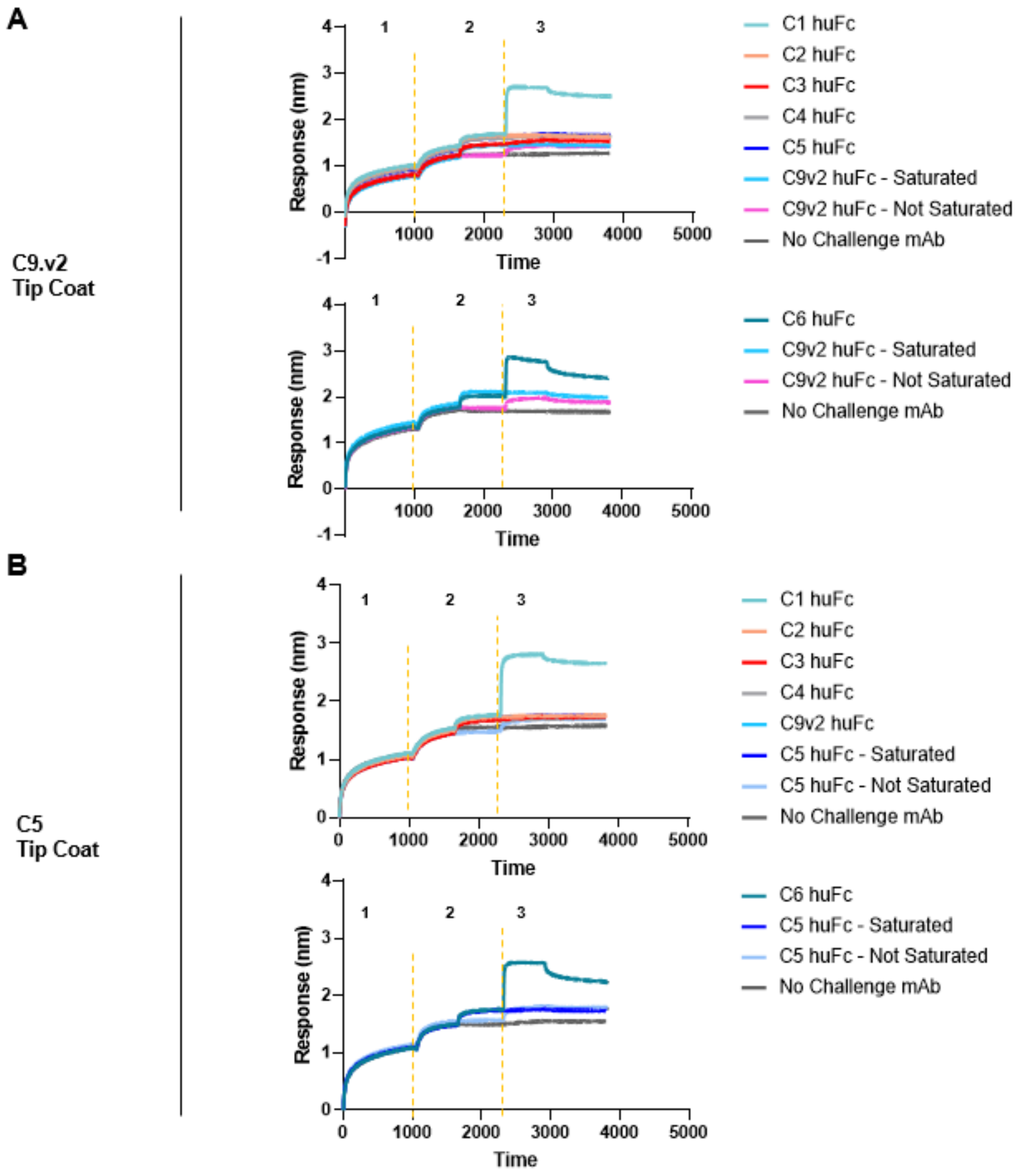


**Supplemental Information 16:** BLI responses used to bin anti-PlGF antibodies into epitope groups using (A) C9.V2 or (B) C5 as the capture antibody. Values shown in main Figure 3 were calculated by subtracting the signal at 500s from the signal at 0s of association (step 3).

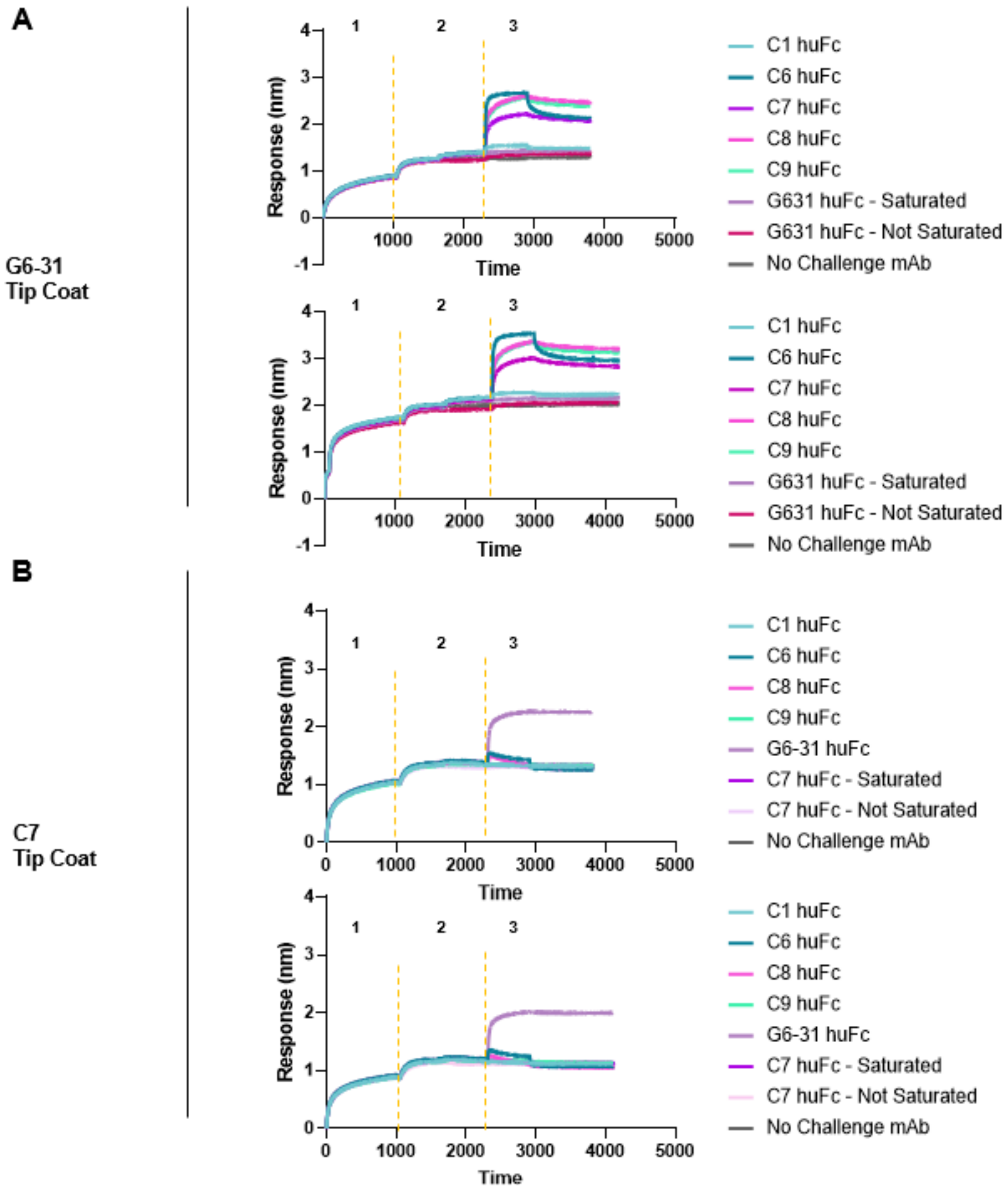


**Supplemental Information 17:** BLI responses used to bin anti-VEGF antibodies into epitope groups using (A) G631 or (B) C7 as the capture antibody. Values shown in main Figure 3 were calculated by subtracting the signal at 500s from the signal at 0s of association (step 3).

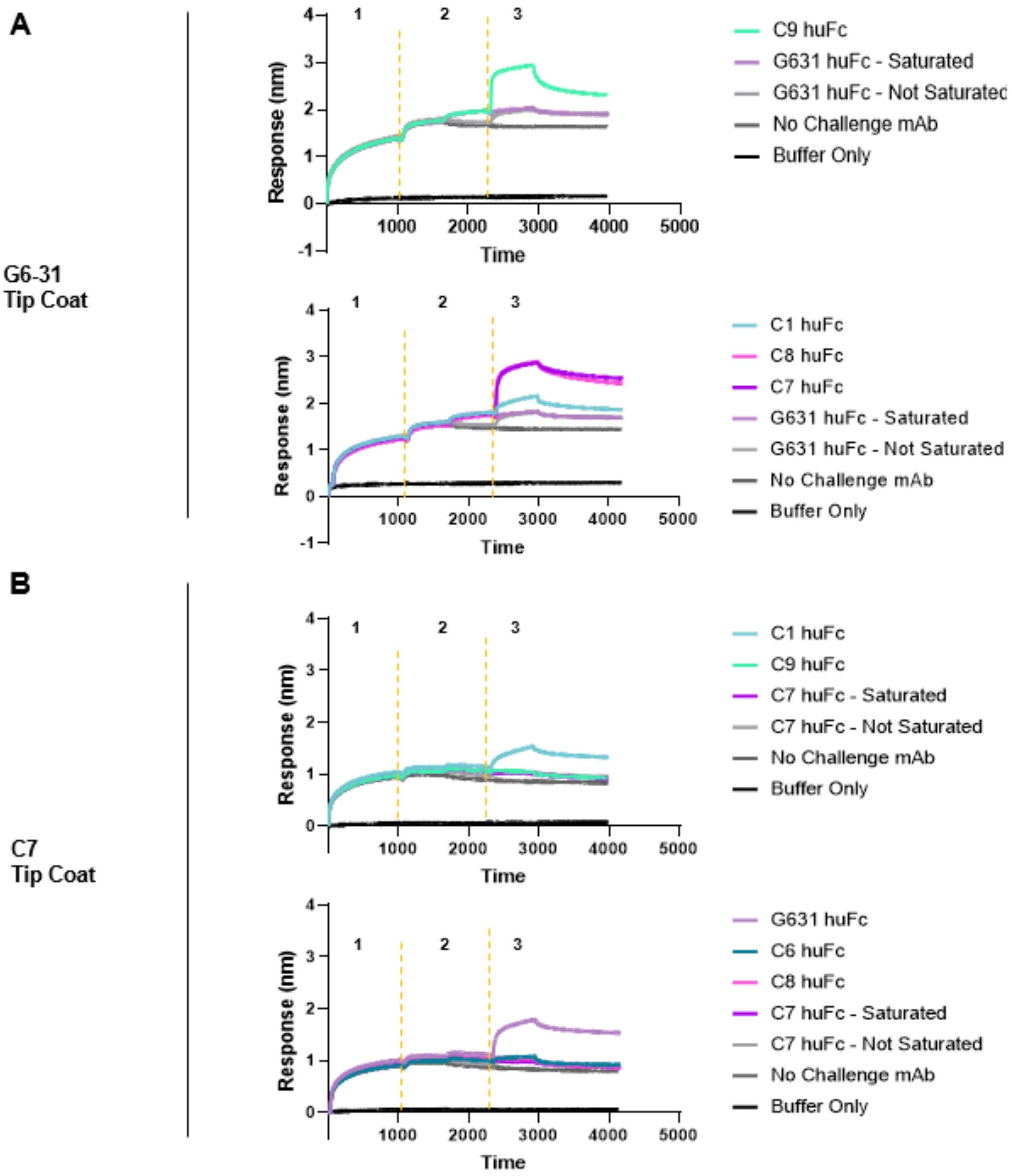


**Supplemental Information 18:** BLI responses used to bin anti-VEGF antibodies into epitope groups using (A) G631 or (B) C7 as the capture antibody. Values shown in main Figure 3 were calculated by subtracting the signal at 500s from the signal at 0s of association (step 3).

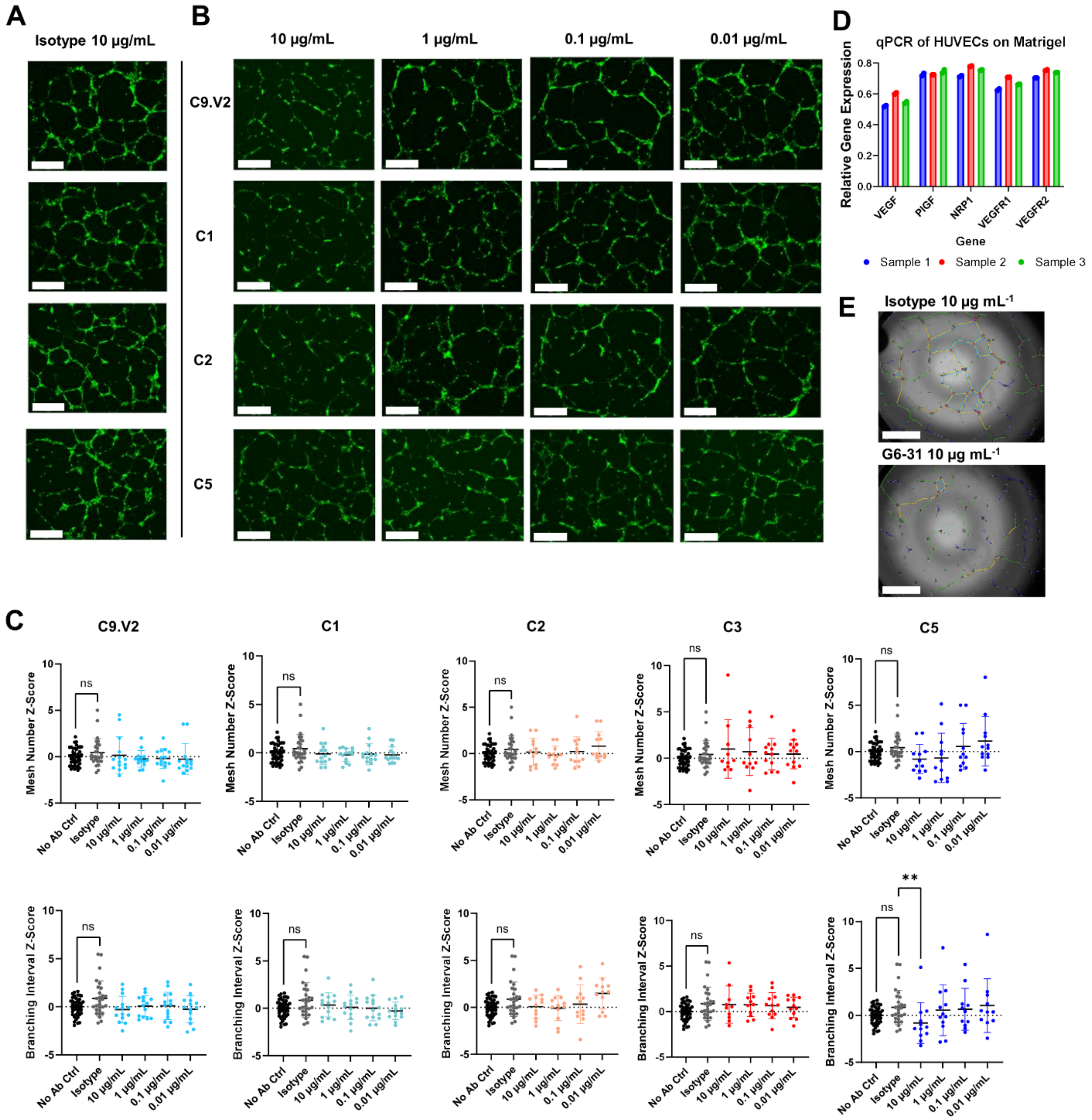


**Supplemental Information 19:** Anti-PlGF antibodies have no effect on HUVEC tube formation. GFP-fluorescent images of HUVEC tube formation depict the isotype dosed at 10 µg/mL (A) and the PlGF Ctrl, C1, C2, and C5 dosed at 10, 1, 0.1, and 0.01 µg/mL (B). (C) Angiogenesis Analyzer ImageJ Plugin measurements of Mesh Number and Branching interval are negligibly different between anti-PlGF antibody conditions and the isotype control. (D) Relative qPCR gene expression of VEGF, PlGF, NRP1, VEGFR1, and VEGFR2 derived from three sets of 16-pooled HUVEC tube formation replicates. (E) Example of the Angiogenesis Analyzer ImageJ plugin measuring tube formation for isotype- and G6-31-treated cells.

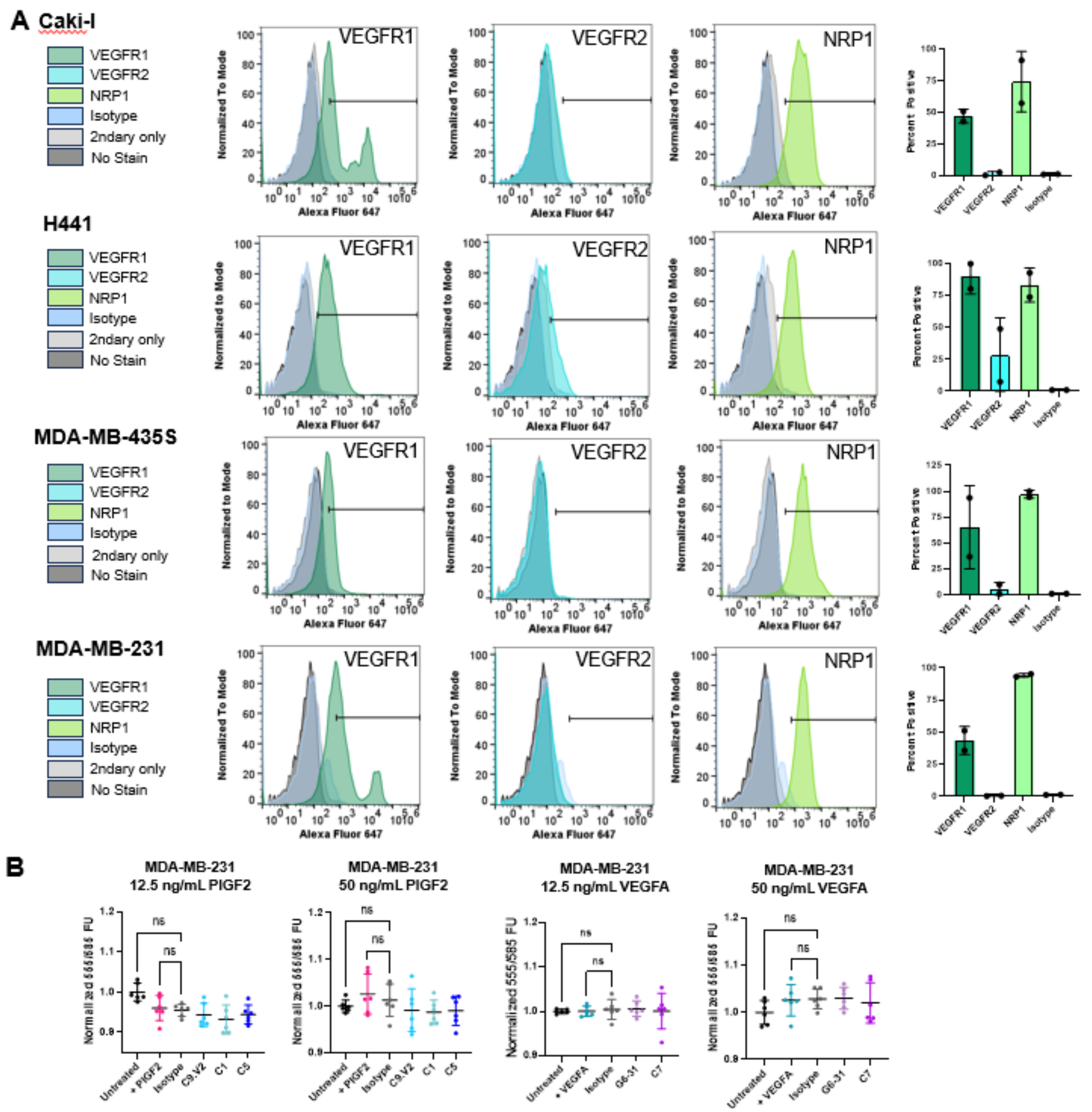


**Supplemental Information 20:** (A) Flow cytometry and percent positive of Caki-I, H441, MDA-MB-435S, and MDA-MB-231 cancer lines stained for VEGFR1, VEGFR2, and NRP1. Percent positive was defined by ~1% overlap with the isotype control staining. (B) Metabolic Inhibition Assay with MDA-MB-231 cells which demonstrates cells were not stimulated by PIGF2 or $VEGFA_{165}$ at 12.5 or 50 ng/mL after starving in serum-free media.

**A.**

| Fab | *AF3 Docking to PlGF2 NRP1 domain* | | *AF3 Docking to VEGFA NRP1 domain* | |
|---|---|---|---|---|
| | ipTM | pTM | ipTM | pTM |
| C1 | 0.81 | 0.86 | N/A | N/A |
| C6 | 0.78 | 0.86 | 0.73 | 0.82 |
| C7 | N/A | N/A | 0.72 | 0.81 |
| Fab | *AF3 Docking to PlGF1 (excludes NRP1 domain)* | | *AF3 Docking to PlGF2 (includes NRP1 domain)* | |
| C9.V2 | *0.42* | *0.55* | 0.38 | 0.51 |
| C5 | *0.44* | *0.58* | 0.5 | 0.61 |

### B. Antibody C1 docked with the PlGF2 heparin binding domain

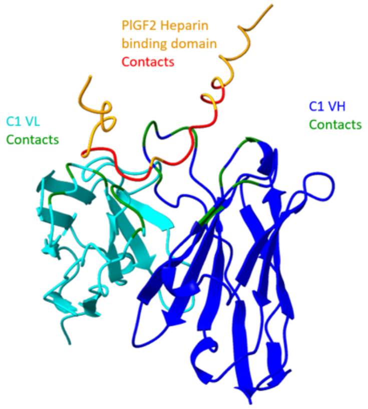


**>C1 VH with predicted contacts in green**

EVQLQQSGAELVRSGASVKLSCTASG**FNIKDYYMH**WVKQRPEQGLEWIG**WIDPEDGDTEYAP**
**KFQG**KATMTADTSSNTAYLQLSSLTSEDTAVYYCNA**PDDDYDSSYAMDY**WGQGTSVTVS

**>C1 VL with predicted contacts in green**

DIVMTQSQKFMSTSVGDRVSVTC**KASQNVGTNVA**WYQQKPGQSPKALIY**SASYRYS**GVPDRF
TGSGSGTDFTLTISNVQSEDLAEYIC**QQYDTFPYT**FGGGTKLEIKR

**>PlGF2 Heparin binding domain with predicted contacts in green**

VH contacts: RRPKGRGKRRREKQRPTDCHLCGDAVPRR

VL contacts: RRPKGRGKRRREKQRPTDCHLCGDAVPRR

### C. Antibody C6 docked with VEGFA heparin binding domain

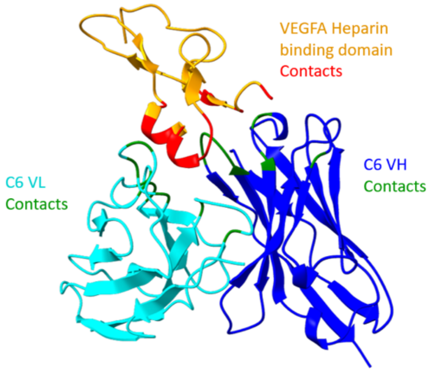


**>C6 VH sequence with predicted contacts in green**

DVKLRESGAELVRSGASVKLSCTASGFNIKDYYMHWVKQRPEQGLEWIGWIDPENGDTEYAPKFQGKATMTADTSSNTAYLQLSSLTSEDTAVYYCNAWGAMDYWGQGTSVTVS

**>C6 VL sequence predicted contacts in green**

DIVMTQSPASLAVSLGQRATISCKASQSVDYDGDSYMNWYQQKPGQPPKLLIYAASNLESGIPARFSGSGSGTDFTLNIHPVEEEDAATYYCQQSYEDPWTFGGGTKLEIKR

**>VEGFA heparin binding domain predicted contacts in green**

$V_H$ contacts: NPCGPCSERRKHLFVQDPQTCKCSCKNTDSRCKARQLELNERTCRCDKPRR

$V_L$ contacts: NPCGPCSERRKHLFVQDPQTCKCSCKNTDSRCKARQLELNERTCRCDKPRR

### D. Antibody C6 docked with PlGF2 heparin binding domain

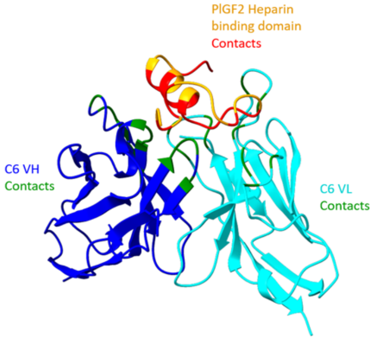


### C6 VH Contacts PlGF2 Heparin Binding Domain

**>C6 VH sequence with predicted contacts in green**

DVKLRESGAELVRSGASVKLSCTASGFNIKDYYMHWVKQRPEQGLEWIGWIDPENGDTEYAPKFQGKATMTADTSSNTAYLQLSSLTSEDTAVYYCNAWGAMDYWGQGTSVTVS

**>C6 VL sequence with predicted contacts in green**

DIVMTQSPASLAVSLGQRATISCKASQSVDYDGDSYMNWYQQKPGQPPKLLIYAASNLESGIPARFSGSGSGTDFTLNIHPVEEEDAATYYCQQSYEDPWTFGGGTKLEIKR

**>PlGF2 heparin binding domain predicted contacts in green**

VH: RRPKGRGKRRREKQRPTDCHLCGDAVPRR

VL: RRPKGRGKRRREKQRPTDCHLCGDAVPRR

### E. Antibody C7 docked with VEGFA heparin binding domain

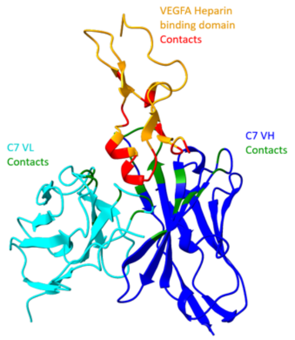


#### C7 VH contacts VEGFA heparin binding domain

##### >C7 VH sequence with predicted contacts in green

QVQLQQSGAELVRSGASVKLSCTASG**FNIKDYYMH**WVKQRPEQGLEWIGW**IDPENGDTEYAPKFQG**KATMTADTSSNTAYLQLSSLTSEDTAVYYCNA**WYYGSSYNYFDY**WGQGTTLTVS

##### >C7 VL sequence with predicted contacts in green

DIVMTQAQKFMSTSVGDRVSVTC**KASQNVGTNVA**WYQQKPGQSPKALI**YSASYRYS**GVPDRFTGSGSGTDFTLTNSNVQSEDLAEYFC**QQYNSYPPT**FGGGTKLEIKR

##### >VEGFA heparin binding domain predicted contacts in green

VH: NPCGPCSERRKHLFVQDPQTCKCSCKNTDSRCKARQLELNERTCRCDKPRR

VL: NPCGPCSERRKHLFVQDPQTCKCSCKNTDSRCKARQLELNERTCRCDKPRR

**Supplemental Figure 21:** AlphaFold3 docking between antibodies and their respective target growth factor subunit, as determined biochemically. **A,** table of AF3 docking scores. pTM score above 0.5 confers the overall predicted fold for the docked complex is likely similar to the true complex. An ipTM score measures the “accuracy of the predicted relative positions of the subunits within the complex.” Values over 0.8 confer high confidence predictions, whereas 0.6-0.8 are in the “gray zone” in which they may or may not be correct. Contacts in the dockings below were determined in ChimeraX by selecting residues with buried solvent-accessible surface area >= 15 Å$^2$.
